# The human ovary exhibits dynamic molecular remodeling in the decades post-menopause

**DOI:** 10.64898/2026.03.26.714635

**Authors:** Mark A. Watson, Bikem Soygur, Christina D. King, Pooja R. Devrukhkar, Simon Melov, Elisheva D. Shanes, Mary Ellen G. Pavone, Francesca E. Duncan, Birgit Schilling

## Abstract

The human ovary is among the first organs to show age-related functional decline, resulting in menopause. Beyond this transition, the postmenopausal ovary is often regarded as quiescent and remains poorly characterized. We analyzed the proteomes of healthy, non-pathological ovaries using mass spectrometry (data-independent acquisition) from 28 postmenopausal women (50-75 years old), stratified into three age groups (50-59, 60-69, ≥70). We quantified 5,812 protein groups and observed progressive age-associated shifts, with 117 proteins significantly altered in the ≥70 vs 50-59 age comparison. Multivariate analysis demonstrated clear separation between 50-59 and ≥70-year-old age cohorts, with protein signatures shifting from RNA/gene-regulatory functions in younger ovaries to metabolic, trafficking, and innate immune/complement pathways in older ovaries. Across differential abundance, multivariate modelling, and covariate-adjusted linear modelling, we identified a convergent set of age-associated proteins that were integrated into the 26-protein **<u>B</u>**uck **<u>P</u>**ostmenopausal **<u>O</u>**vary **<u>M</u>**olecular **<u>S</u>**ignature (BuckPOMS), a consensus molecular signature capturing progressive extracellular matrix remodeling, inflammatory signaling, and loss of structural and keratin-associated proteins with age. Representative BuckPOMS proteins, including the secreted matrisome proteins WNT4 and Fibromodulin (FMOD), were validated by immunohistochemistry. Pathway enrichment further identified an increase in inflammatory and matrisome pathways, and increased abundance of damage-associated secretory factors decades following menopause. These data fundamentally shift the notion of the postmenopausal ovary as an inert organ and instead demonstrate active and continuous molecular remodeling that has potential relevance to tissue signaling and implications for women’s health.

## Introduction

The ovary is recognized as one of the most rapidly aging organs in the human body, with functional decline evident decades before overt aging of other organs is observed (M.J. Faddy, Gosden, Gougeon, Richardson, & Nelson, 1992; Hansen et al., 2008; Knowlton, Craig, Zavy, & Hansen, 2014). Women are born with a fixed number of ovarian follicles, or the ovarian reserve, which contribute to fertility and endocrine function. The ovarian reserve, which depletes with age, dictates reproductive lifespan and determines the timing of menopause (Hansen et al., 2008; Knowlton et al., 2014; Mondal, Tcherniak, & Kolomeisky, 2025; Nikolaou & Templeton, 2004; Shadyab et al., 2017). Menopause, defined as the permanent cessation of menstruation for at least 12 consecutive months, typically occurs around age 50 and marks the end of ovarian reproductive function (Ambikairajah, Walsh, & Cherbuin, 2022; Harlow et al., 2012; Mondal et al., 2025). Menopause onset is linked to increased risks of ovarian cancer (Abulajiang, Liu, Wang, Abulai, & Wu, 2025; Kim et al., 2026) and chronic conditions, including cardiovascular disease (El Khoudary et al., 2020; Hu et al., 1999; Raj et al., 2023), osteoporosis (Ji & Yu, 2015; Yong & Logan, 2021), and neurodegeneration (Briceno Silva et al., 2024; Mosconi et al., 2021; Y. Wang, Mishra, & Brinton, 2020; Wood Alexander et al., 2025), highlighting the ovary’s disproportionate influence on systemic aging in women and overall health span. Due to medical and health advances, many women can expect to live decades in a postmenopausal state. Despite our increased longevity, we know surprisingly very little about the biology and function of the postmenopausal ovary.

The longstanding view of the postmenopausal ovary is that it is largely composed of hormonally inactive fibrous connective tissue (Broekmans, Soules, & Fauser, 2009; Clement, 1987; M. J. Faddy & Gosden, 1996; Laszczynska, Brodowska, Starczewski, Masiuk, & Brodowski, 2008; Motta, Makabe, & Nottola, 1997; Richardson, Senikas, & Nelson, 1987; Tepper, Zalel, Markov, Cohen, & Beyth, 1995). Classical histological profiling typically describes the postmenopausal ovary as a reduced-volume, fibrotic organ with follicle depletion, cortical thinning, loss of cortico-medullary distinction, epithelial invaginations (inclusion cysts), and abundant corpora albicantia. In fact, ovarian fibrosis, characterized by fibroblast expansion and extracellular matrix (ECM) accumulation, is a key contributor to age-related ovarian dysfunction (Amargant et al., 2020; Briley et al., 2016; Gu, Wang, & Yu, 2024; Landry, Vaishnav, & Vanderhyden, 2020; Umehara et al., 2022). However, emerging data suggest that the postmenopausal ovary may not be as quiescent as previously thought. Even after menopause, the ovary can retain endocrine and paracrine activity, including androgen production and local signaling, with potential systemic effects (Fogle, Stanczyk, Zhang, & Paulson, 2007; Judd, Lucas, & Yen, 1974; Rinaudo & Strauss, 2004).

At the molecular level, single-nucleus, single-cell, and spatial transcriptomic atlases of human ovaries are beginning to reveal coordinated age-related changes in signaling, metabolism, and stress responses across multiple cell types, echoing aging trajectories observed in other organs (P. R. Devrukhkar et al., 2025; Fan et al., 2019; Jin et al., 2025; Jones et al., 2024; Lengyel et al., 2022; Senchyna et al., 2026; Watson et al., 2025; Wu et al., 2024). However, many studies have focused on ovarian tissue from pre- or peri-menopausal women, with limited sample sizes. As a result, the molecular state of the “aged, normal” human ovary across the decades following reproductive cessation remains underexplored. Similar to transcriptomic studies, existing proteomic studies of the human ovary remain few and are typically limited by small cohorts, with a primary focus on premenopausal tissue or the early menopausal transition (P. R. Devrukhkar et al., 2025; Emna Ouni et al., 2022; E. Ouni, Vertommen, Chiti, Dolmans, & Amorim, 2019; Yi et al., 2018; Zhang et al., 2014). Consequently, we lack a proteome map of the human postmenopausal ovary. To advance the field, we performed data-independent acquisition mass spectrometry using parallel accumulation-serial fragmentation (diaPASEF) (Meier et al., 2020) proteomics on healthy human ovaries from 28 participants (aged 50-75 years) spanning the menopausal transition and over two decades thereafter. Across 5,812 quantified protein groups, we observed progressive age-associated shifts in the ovarian proteome. We combined differential relative abundance analysis, multivariate partial least squares-discriminant analysis (PLS-DA), and covariate-adjusted continuous-age modeling with pathway- and matrisome-focused enrichment analyses to define age-associated proteomic signatures in the postmenopausal ovary. These complementary analytical approaches were integrated to develop the 26-protein Buck Postmenopausal Ovary Molecular Signature (BuckPOMS), a consensus molecular signature of postmenopausal ovarian aging. Finally, we examined whether age-associated proteomic remodeling was enriched for proteins previously implicated in ovarian cancer and for proteins identified in published ovarian cellular senescence signatures to determine whether postmenopausal ovarian aging converges with senescence-adjacent extracellular matrix (ECM) remodeling programs. Together, these analyses reveal that the postmenopausal ovary continues to undergo active molecular remodeling long after reproductive cessation, characterized by shifts toward inflammatory, complement, and matrisome-associated pathways as well as increased damage-associated secretory factors. This ongoing molecular remodeling that occurs in the postmenopausal ovary challenges the notion that the ovary is an innocent bystander in the post-reproductive period and instead suggests a potentially more active role in systemic aging and chronic disease risk in women.

## Results

### Pairwise differential proteomic analysis reveals age-associated remodeling of the postmenopausal ovary

To identify protein signatures of “healthy aging” in the human postmenopausal ovary, we collected a set of non-pathological ovarian tissue from women aged 50-75 years (n = 28) and applied a comprehensive data-independent acquisition (DIA) (Bruderer et al., 2017; Collins et al., 2017; Gillet et al., 2012) mass spectrometry workflow to quantify thousands of proteins in an unbiased manner (Figure 1a & Data S1). Specifically, proteomics analysis was performed on non-pathological ovarian tissue from 28 postmenopausal participants. These tissues were subjected to tissue lysis, proteolytic digestion of proteins, followed by data-independent acquisition mass spectrometry using parallel accumulation-serial fragmentation (diaPASEF) (Meier et al., 2020); Figure 1a). In this approach, the mass spectrometer systematically measures and quantifies peptides across all samples in an unbiased way, and relative protein abundance was quantified across all participants. Participants were aged 50-75 years and stratified into three groups: 50-59 (n = 11), 60-69 (n = 10), and ≥70 years (n = 7; Figure 1b & Data S1). Although only ovaries with normal histological pathology, as assessed by a board-certified gynecological pathologist, were included, participants underwent oophorectomy for a range of clinical indications, including benign ovarian conditions (Figure 1c & Data S1). Endometrial pathology, including endometriosis, was the most common indication. We also performed permutation testing to assess whether any surgical indication was associated with age group. No individual surgical indication remained significantly associated with age after Benjamini-Hochberg multiple-testing correction (Data S1), and endometrial pathology was not enriched in the ≥70-year cohort. Self-reported race and ethnicity showed that most participants identified as white (60.7%; Figure 1d & Data S1).

**Figure 1.**
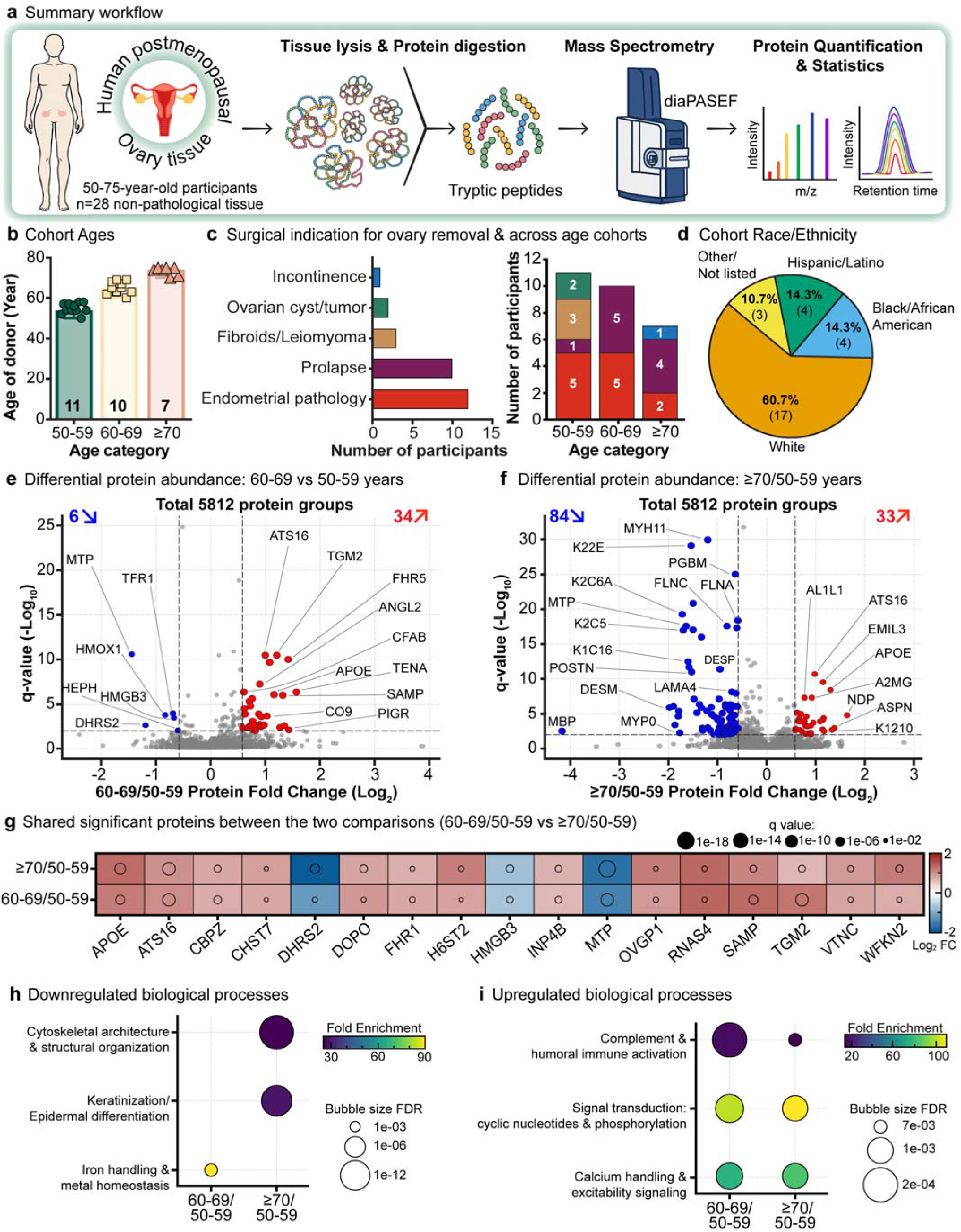
Pairwise differential proteomic analysis across postmenopausal ovarian age cohorts. (**a**) Overview of the workflow for the proteomic evaluation of postmenopausal ovaries. Non-pathological postmenopausal ovarian tissue from women aged 50-75 years was collected, digested to tryptic peptides, and analyzed by data-independent acquisition mass spectrometry using parallel accumulation-serial fragmentation (diaPASEF), followed by protein quantification. (**b**) Participant age distribution and age-group stratification; the number of participants and age range in years for each group are shown (50-59, n = 11; 60-69, n = 10; ≥70, n = 7). (**c**) Primary surgical indications leading to ovary removal and distribution of surgical indications across the three postmenopausal age cohorts. (**d**) Participant distribution of self-reported race and ethnicity across the cohort. (**e-f**) Differential protein abundance between age cohorts. Volcano plots summarize pairwise comparisons using thresholds of log_2_ fold change (log2 FC) >0.58 (∼1.5-fold) and q-value <0.01. (**e**) ovaries from 60-69 vs 50-59-year-old. (**f**) ovaries from ≥70 vs 50-59-year-old. Proteins increased are shown in red, decreased proteins in blue, and proteins not meeting thresholds in grey. (**g**) A heatmap of shared significantly altered proteins (log2 FC >0.58, q-value <0.01) between 60-69 vs 50-59 and ≥70 vs 50-59-year-old age comparisons. Color indicates log2 FC, while the circle size indicates q-value. (**h-i**) Gene Ontology (GO) Biological Process enrichment of significantly altered proteins. (**h**) Enriched biological processes among downregulated proteins and (**i**) enriched processes among upregulated proteins across the age comparisons.

Across all 28 ovarian tissue samples, proteomic analysis yielded 5,812 quantified protein groups with at least 2 unique peptides (Data S2-3). A protein group is a collection of proteins, including isoforms and proteoforms, that share the same identified peptide sequences and therefore cannot be distinguished based on the observed peptides (Nesvizhskii & Aebersold, 2005). We then performed pairwise age-group comparisons with strict statistical thresholds (Log_2_FC >0.58, q <0.01) (Data S4-5). We compared ovarian proteomes from women aged 60-69 to those from women 50-59-years-old and observed that 6 protein groups significantly decreased and 34 increased in the 60-69-year age group (Figure 1e & Data S4). These changes point to a more inflammatory and fibrotic milieu in the ovaries of 60-69-year-old individuals, reflected by higher levels of complement and humoral immune factors, including Complement Factor H-related Protein 1 (FHR1), Complement Factor H-related protein 5 (FHR5), Complement Factor B (CFAB), Complement Component C9 (C9), and Clusterin (CLU). In parallel, matrix/scavenger proteins were also elevated, including A disintegrin and metalloproteinase with thrombospondin motifs 16 (ATS16), Protein-glutamine gamma-glutamyltransferase 2 (TGM2), and Vitronectin (VTNC) (Figure 1e & Data S4).

We then compared ovarian proteomes from women aged ≥ 70 years with those from women aged 50-59 years and observed that 84 protein groups significantly decreased, and 33 significantly increased in the ≥70-year-old age group (Figure 1f & Data S5). Together, these changes suggest a shift from preserved smooth muscle/epithelial architecture toward a more remodeled, secretory tissue state in very late postmenopause. This is reflected by depletion of structural and cytoskeletal proteins such as Myosin-11 (MYH11), Filamin-A and C (FLNA/FLNC), Dystrophin (DMD), and multiple keratins (K22E, K2C1, K2C6A, K1C14), alongside enrichment of extracellular remodeling and damage-associated secretory proteins including ATS16, EMILIN-3 (EMIL3), Norrin (NDP), Alpha-2-macroglobulin (A2M), and TGM2 (Figure 1f & Data S5).

A core set of seventeen proteins was significantly altered in both pairwise age comparisons (60-69 vs 50-59-year-old and ≥70 vs 50-59-year-old), and these shared proteins indicate overlapping biology across the postmenopausal decades (Figure 1g & Data S4-5). Many of these proteins map to complement/innate immunity and matrisome/extracellular matrix (ECM) remodeling, including Complement factor H-related protein 1 (FHR1), A disintegrin and metalloproteinase with thrombospondin motifs 16 (ATS16), Vitronectin (VTNC), Protein-glutamine gamma-glutamyltransferase 2 (TGM2), Carboxypeptidase Z (CBPZ), Serum amyloid P-component (SAMP), and WAP, Kazal, immunoglobulin, Kunitz, and NTR domain-containing protein (WFKN2), all of which are consistently higher in the older groups (Figure 1g). The overlap also implicates lipid handling, secreted glycoproteins, and sulfated glycosaminoglycan metabolism, as reflected by Apolipoprotein E (APOE), Microsomal triglyceride transfer protein large subunit (MTP), Dehydrogenase/reductase SDR family member 2, mitochondrial (DHRS2), Heparan-sulfate 6-O-sulfotransferase 2 (HS6ST2), and Oviduct-specific glycoprotein (OVGP1) (Figure 1g).

These protein signatures are mirrored at the pathway level. Proteins that are significantly downregulated map to cytoskeletal architecture/structural organization and keratinization/epidermal differentiation in the ≥70 vs 50-59-year-old comparison, with iron handling and metal homeostasis reduced by 60-69 years (Figure 1h & Data S6). In contrast, upregulated proteins are enriched for complement and humoral immune activation in both age cohort comparisons, together with cyclic nucleotide/phosphorylation signaling and calcium handling/excitability signaling, indicating a shift from a structurally maintained, epithelial/keratin-rich ovary toward one dominated by inflammatory and Ca²⁺-driven signaling with advancing postmenopausal age (Figure 1i & Data S6).

### Multivariate proteomic analysis identifies protein signatures that distinguish younger and older postmenopausal ovaries

While pairwise differential analysis identified proteins that changed between age groups, we next used multivariate proteomic analysis to determine whether the overall ovarian proteome also separated by postmenopausal age and to identify the proteins that best drove this separation. A three-class partial least squares-discriminant analysis (PLS-DA) model including all age groups (50-59, 60-69, and ≥70-years-old) showed only modest separation (Figure S1a & Data S7-8). The 60-69 and ≥70-year-old groups largely overlapped, whereas the 50-59-year-old group separated slightly from the older cohorts (Figure S1a & Data S7-8). This limited discrimination was reflected by a mean 5-fold cross-validated accuracy of 0.44. Pairwise PLS-DA models performed better and separated the groups from each other: the 50-59 vs 60-69-year-old comparison achieved a modest separation reflected by the mean accuracy of 0.52 (Figure 2a & Data S7-8), whereas the 50-59 vs ≥ 70-year-old comparison had a greater separation, which reached an accuracy score of 0.70 (Figure 2b & Data S7-8), with Latent Variable 1 (LV1) capturing the major age-related gradient in both cases. To further assess model performance, we generated receiver operating characteristic (ROC) curves from cross-validated predictions for the two pairwise models (Figure 2c). In line with the clearer separation seen in the PLS-DA LV1/LV2 plots, the 50-59 vs ≥70 year old comparison discriminated age groups substantially better (AUC ≈ 0.73) than the 50-59 vs 60-69-year-old comparison (AUC ≈ 0.52), indicating that larger age differences were accompanied by a stronger and more coherent shift in the ovarian proteomes (Figure 2c).

**Figure 2.**
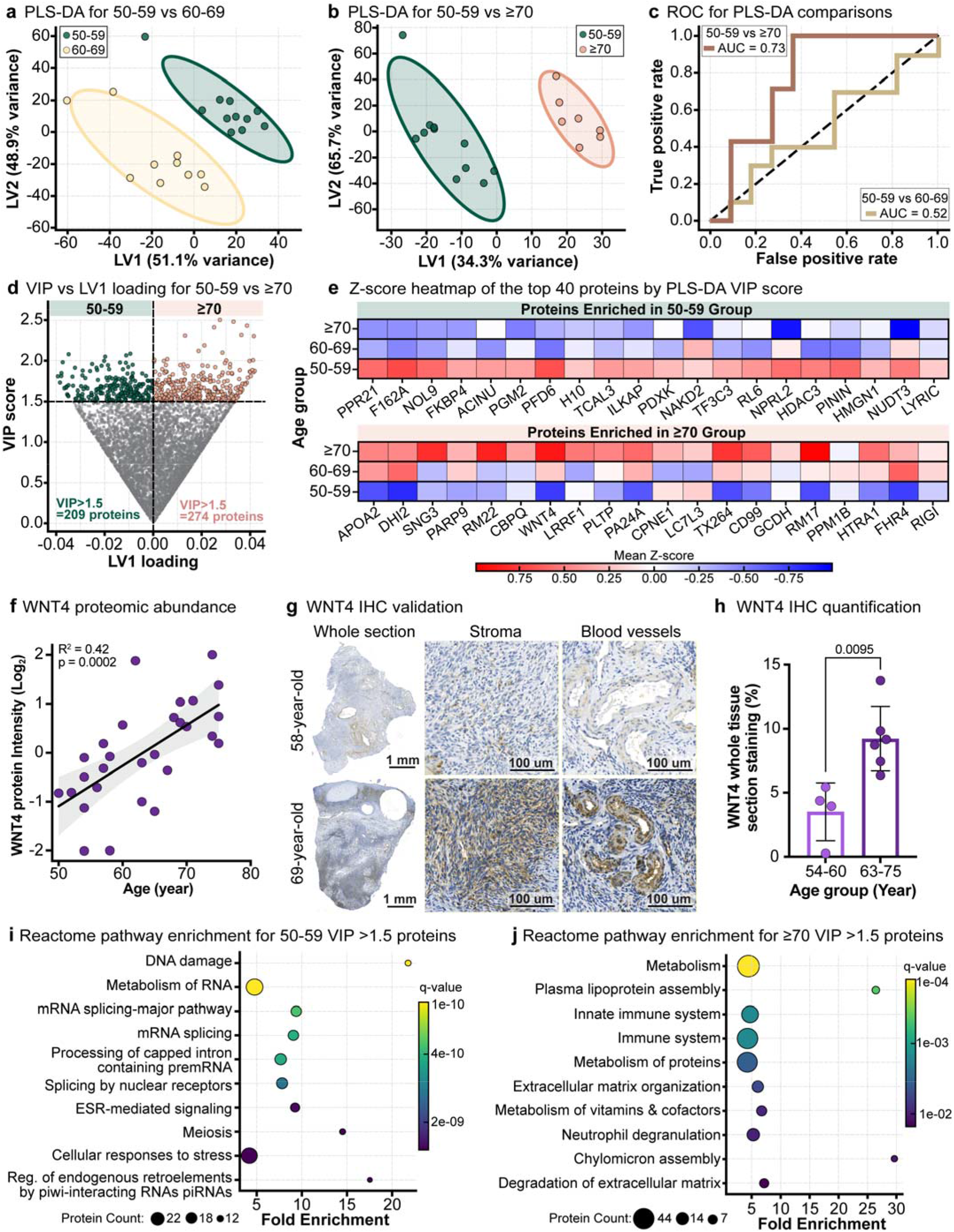
Multivariate proteomic profiling identifies VIP-defined protein signatures that distinguish younger and older postmenopausal ovaries. (**a-b**) Pairwise partial least squares-discriminant analysis (PLS-DA) comparing (**a**) ovaries from women aged 50-59 vs 60-69 years and (**b**) ovaries from 50-59 vs ≥70 years. Score plots show the first two latent variables (LV1 and LV2); each point represents a participant, and ellipses indicate group dispersion in LV space. (**c**) Cross-validated receiver operating characteristic (ROC) curve for the 50-59 vs ≥70-year-old PLS-DA model. Model performance is summarized by the area under the curve (AUC). (**d**) Variable importance in projection (VIP) score vs LV1 loading for the 50-59 vs ≥70-year-old PLS-DA model. Proteins with VIP ≤1.5 are shown in grey. Proteins with VIP >1.5 are colored by direction: higher abundance in ≥70-year-olds is shown in pink, and higher abundance in 50-59-year-olds is shown in green. (**e**) Heatmap of z-scored log_2_ protein abundance for the top VIP-selected proteins (20 ≥70-year-old-associated and 20 50-59-year-old-associated) across all participants, highlighting coordinated age-associated shifts in the multivariate signature. (**f**) Association of WNT4 protein abundance with age. Linear regression is shown with 95% confidence interval (shaded); R² and p-value are indicated. (**g**) Representative WNT4 immunohistochemistry (IHC) in a younger (58-year-old) and older (69-year-old) participant. Whole-section images are shown alongside higher-magnification views of stroma and blood vessels. Scale bars are shown in the lower right. (**h**) Quantification of WNT4 IHC staining across 10 participants. Unpaired two-sided t-test; p-value shown. Reactome pathway gene set enrichment analysis using proteins ranked by VIP >1.5 and signed LV1 loading. (**i**) Pathways enriched in 50-59-year-old ovaries and (**j**) pathways enriched in ≥70-year-old ovaries.

To identify proteins that most strongly drove separation between ovaries from women aged 50-59 and ≥70 years in the PLS-DA model, we examined variable importance in projection (VIP) scores and LV1 loadings. The VIP score summarizes each protein’s contribution to group discrimination, whereas the LV1 loadings provide directionality, indicating whether proteins were relatively higher in 50-59- or ≥70-year-old ovaries. Although VIP >1 is commonly used as a standard threshold, we applied a more stringent cutoff of VIP >1.5 to focus on the strongest contributors (Akarachantachote, Chadcham, & Saithanu, 2014; Chong & Jun, 2005). In total, 209 proteins with VIP >1.5 and negative LV1 loadings were associated with the 50-59-year age cohort, whereas 274 proteins with VIP >1.5 and positive LV1 loadings were associated with the ≥70-year age cohort (Figure 2d & Data S9). A heatmap of the top 40 VIP-selected proteins (20 proteins distinguishing 50-59-year-olds, and 20 proteins distinguishing ≥70-year-olds) showed a clear bidirectional pattern: proteins enriched in 50-59-year-olds were consistently higher in the 50-59-year-old group and lower in the ≥70-year-old group (Figure 2e). However, proteins enriched for ≥70-year-olds showed the opposite trend, with the 60-69-year age group generally displaying intermediate values (Figure 2e). In 50-59-year-old ovaries, the multivariate signature was dominated by proteostasis and chromatin/gene-regulation machinery. Key contributors included Peptidyl-prolyl cis-trans isomerase FKBP4 (FKBP4) and Prefoldin subunit 6 (PFD6) (proteostasis/protein folding), together with non-histone chromosomal protein HMG-14 (HMGN1) and Histone deacetylase 3 (HDAC3) (chromatin/gene regulation) (Figure 2e). In contrast, the protein signature from the≥ 70-year-old ovaries shifted toward lipid metabolism and innate immunity. Proteins enriched in ≥70-year-olds included Apolipoprotein A-II (APOA2), Phospholipid transfer protein (PLTP), and 11-beta-hydroxysteroid dehydrogenase type 2 (DHI2) (lipid handling/metabolism), as well as immune-associated proteins such as Protein mono-ADP-ribosyltransferase PARP9 (PARP9), Complement factor H-related protein 4 (FHR4), and Antiviral innate immune response receptor RIGI (RIGI) (Figure 2e).

WNT4 emerged as a particularly interesting candidate because it is a secreted signaling ligand with established roles in ovarian development and stromal organization (Boyer et al., 2010; Prunskaite-Hyyrylainen et al., 2016; Vainio, Heikkila, Kispert, Chin, & McMahon, 1999). Its identification, therefore, raised the possibility that the aging postmenopausal ovary retains not only remodeling features but also active signaling capacity. WNT4 abundance progressively increased with age across ovaries from women aged 50-59, 60-69, and ≥70 years (Figure 2e), and it was identified as a significant candidate in the pairwise differential analysis (Figure 2f & Data S5). In the proteomic dataset, WNT4 abundance was positively associated with age (R² = 0.42, p = 0.0002) (Data S7). Immunohistochemical (IHC) staining of WNT4 in ovarian tissue sections from 10 participants confirmed increased abundance, particularly in the ovarian stroma and blood vessels of older participants (Figure 2g). Quantification of WNT4-positive staining further showed higher abundance in the 63-75-year-old group (Figure 2h) and a positive correlation with continuous age (R² = 0.41, p = 0.06; Figure S1b).

Reactome pathway enrichment of proteins with VIP>1.5 revealed distinct age-stratified pathway signatures. In the 50-59-year group, enriched pathways were dominated by RNA metabolism and mRNA processing (including capped intron-containing pre-mRNA processing and mRNA splicing), alongside gene expression and transcription, epigenetic regulation, and broad cellular stress/stimulus response terms, consistent with a gene-regulation and RNA-processing program in younger postmenopausal ovaries (Figure 2i & Data S10). In contrast, proteins driving the ≥70-year-old multivariate signature are heavily enriched for metabolic and immune/innate immune pathways, with broad metabolism and metabolism of proteins among the strongest hits. We also observed in the ≥70-year-olds, prominent enrichment of lipoprotein/lipid handling (plasma lipoprotein and chylomicron assembly) and extracellular matrix organization/degradation, together with neutrophil degranulation, pointing to a coordinated shift toward metabolic remodeling, lipid transport, innate immunity, and ECM turnover in the oldest ovaries (Figure 2j Data S10).

### Continuous-age regression reveals proteins that progressively change across postmenopausal aging

After observing age-group differences in the ovarian proteome, we next asked which proteins changed progressively with age across the full cohort, while accounting for potential confounding by body mass index (BMI) and self-reported race/ethnicity. We therefore performed covariate-adjusted regression, modeling age as a continuous variable while controlling for BMI and self-reported race/ethnicity. We compared a linear model (abundance ∼ Age + BMI + Race/ethnicity) with a quadratic model (abundance ∼ Age + Age² + BMI + Race/ethnicity) (Figure 3a & Data S11-13). Model comparison using complementary metrics: Akaike Information Criterion (AIC), Bayesian Information Criterion (BIC), and an F-test for nested models, consistently favored the linear specification, indicating that inclusion of an Age² term did not improve fit for most proteins (AIC: 3,852/5,806 proteins; BIC: 4,718/5,806 proteins) (Figure S2a-d & Data S13). We therefore based the subsequent age-association analyses on the linear regression model.

**Figure 3.**
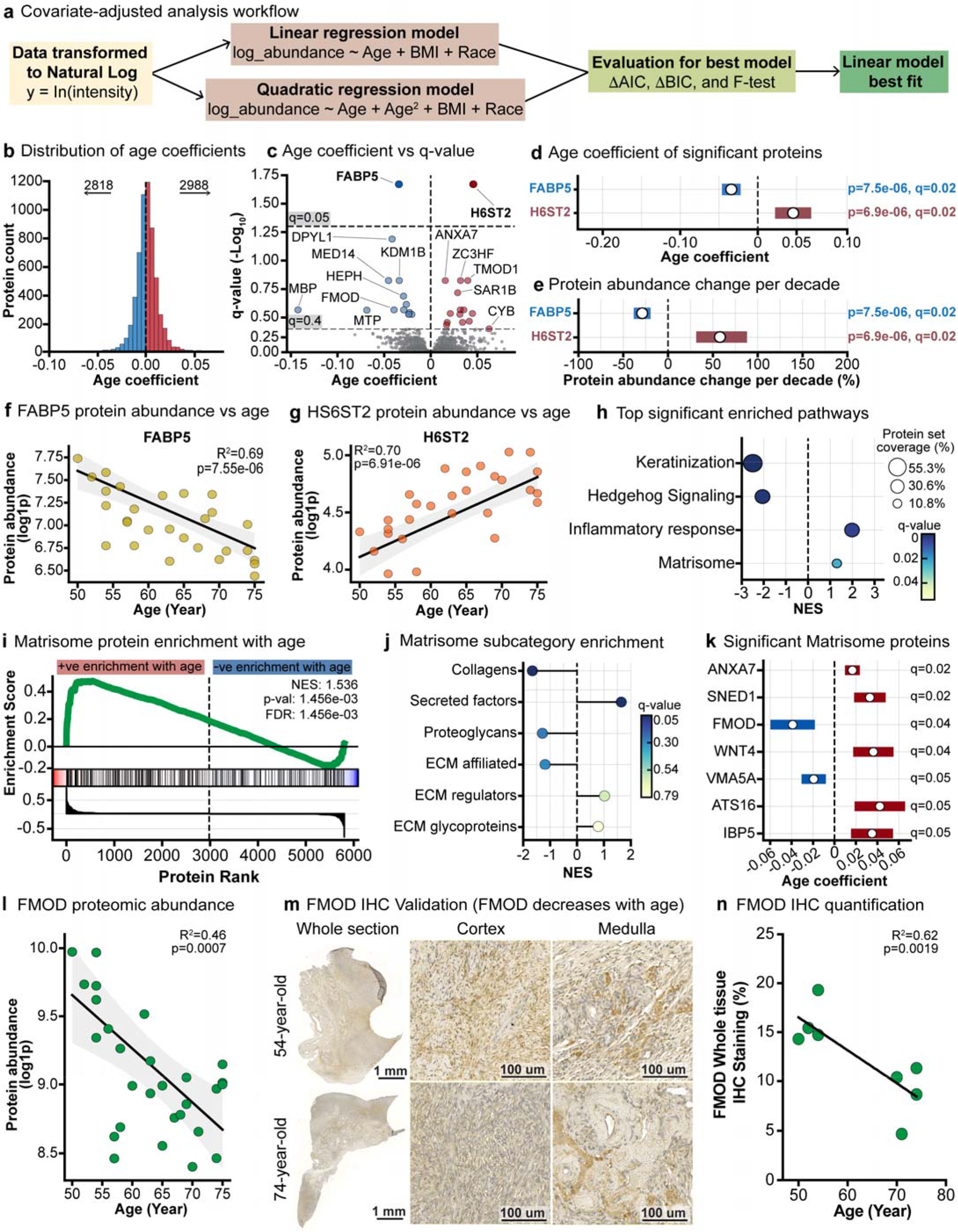
Covariate-adjusted regression identifies age-associated proteins and pathways in the postmenopausal ovary. (**a**) Overview of the covariate-adjusted modeling workflow. Model comparison indicated that a linear age term provided the best fit; the linear model was used for all subsequent analyses. AIC; Akaike Information Criterion, BIC; Bayesian Information Criterion. (**b**) Distribution of age coefficients (β_age) across quantified proteins. Negative coefficients (blue) indicate decreased abundance with age, and positive coefficients (red) indicate increased abundance with age. (**c**) Volcano plot of β_age versus statistical significance (q-value) for all proteins. The dashed line indicates the significance threshold (q <0.05). The two significant proteins are highlighted, along with the top-ranked candidate proteins (q <0.4). (**d**) Forest plot showing β_age estimates (±95% CI) for the two significant age-associated proteins (q <0.05). (**e**) Forest plot showing the corresponding percent change in protein abundance per decade, derived from β_age, for the two significant proteins (q <0.05). (**f**) Association of FABP5 ln(1 + abundance) with age across participants. Linear regression is shown with 95% confidence interval (shaded); R² and p-value are indicated. (**g**) Association of HS6ST2 ln(1 + abundance) with age across participants. Linear regression is shown with 95% confidence interval (shaded); R² and p-value are indicated. (**h**) Gene set enrichment analysis (GSEA) using MSigDB Reactome and Hallmark gene sets was performed on age association scores derived from the covariate-adjusted linear model fit to ln(1 + abundance). The most significantly enriched pathways are shown (q-values indicated). (**i**) Enrichment analysis using MatrisomeDB, reporting normalized enrichment score (NES) and FDR (q-value) to assess age-associated enrichment of matrisome proteins. (**j**) Enrichment of MatrisomeDB subcategories to identify which matrisome components drive the overall signal. (**k**) Targeted covariate-adjusted linear regression restricted to MatrisomeDB proteins, highlighting the most age-associated matrisome candidates. (**l**) Association of FMOD ln(1 + abundance) with age across participants. Linear regression is shown with 95% confidence interval (shaded); R² and p-value are indicated. (**m**) Representative FMOD immunohistochemistry (IHC) in a younger (54-year-old) and older (74-year-old) participant. Whole-section images are shown alongside higher-magnification views of cortical and medullary regions. Scale bars are shown in the lower right. (**n**) Quantification of FMOD IHC staining across 8 participants. Linear regression fit is shown with R² and p-value indicated. Note: log1p(abundance) corresponds to ln(1 + abundance) and was used for panels **f**, **g**, and **l**.

The linear model yielded an age-coefficient (β_age) distribution in which 2,818 proteins decreased with age (β_age <0) and 2,988 increased with age (β_age >0) (Figure 3b & Data S12). Plotting β_age against q-value identified two significant age-associated proteins (q <0.05): Fatty acid-binding protein 5 (FABP5) decreased with age (β_age = −0.034, q = 0.02), and Heparan-sulfate 6-O-sulfotransferase 2 (H6ST2) increased with age (β_age = 0.046, q = 0.02) (Figure 3c-d, 3f-g, & Data S12). For interpretability, we converted β_age into the estimated percent change in protein abundance per decade and plotted these effect sizes for the significant proteins (Figure 3e and Figure S3b). FABP5 decreased in abundance by ∼-29% per decade, and H6ST2 increased in abundance by ∼58% per decade (Figure 3e and Figure S3b). Relaxing the threshold to q < 0.4 revealed an additional 28 proteins that are highlighted in age group comparison analyses, including WNT4, FMOD, ATS16, and IBP5 (Figure 3c, Figure S3a-b, & Data S12). When examining the top 30 age-associated proteins ranked by q-value, they group into distinct functional clusters, including proteins involved in extracellular matrix organization and remodeling (Figure S4a), lipid metabolism and secretory trafficking (Figure S4b), gene regulation and cell cycle control (Figure S4c), and cytoskeletal dynamics and membrane trafficking (Figure S4d).

### Pathway enrichment reveals gain of inflammatory and matrisome programs and loss of structural and keratin-associated programs with age

Using the age coefficient from the linear model as a ranking metric, we performed gene set enrichment analysis (GSEA) with MSigDB Reactome (Matthews et al., 2009), Hallmark (Liberzon et al., 2015), and MatrisomeDB (Naba et al., 2012; Shao et al., 2023) gene sets to identify age-associated pathways (Figure 3h, Figure S5a & Data S14). Significant pathways (FDR < 0.05) included downregulated keratinization (Normalized enrichment score (NES) = -2.491, FDR < 0.0001; Figure S5b & & Data S14) and Hedgehog signaling (NES = –2.054, FDR = 0.00366; Figure S5c & Data S14), as well as upregulated inflammatory response (NES = 1.984, FDR = 0.009; Figure S5d & Data S14) and matrisome-related pathways (NES = 1.54, FDR = 0.0014; Figure 3i & Data S14). Because matrisome remodeling is such a prominent feature of ovarian aging, we used MatrisomeDB subcategories to examine which classes of matrisome proteins were most strongly associated with age (Figure 3j & Data S15). This analysis revealed significant negative enrichment of collagens and positive enrichment of secreted factors with age (p < 0.05), consistent with our earlier findings of age-related shifts in the ovarian extracellular matrix (Figure 2j). A targeted covariate-adjusted linear regression restricted to MatrisomeDB proteins highlighted the most age-associated matrisome candidates sorted by q-value (Figure 3k & Data S16). This notably included WNT4, which we independently validated as a candidate marker of aged ovaries (Figure 2g-h).

Fibromodulin (FMOD) emerged as a particularly compelling matrisome candidate because it was identified in both the pairwise differential analysis (Data S5) and the matrisome-targeted continuous-age regression (Figure 3k & Data S16), and because its known biological functions directly align with the extracellular remodeling and fibrotic signatures observed in older ovaries. FMOD was significantly downregulated with age in the linear model (β_age = -0.039, q = 0.04; Figure 3k & Data S16). FMOD is a small leucine-rich extracellular matrix proteoglycan that organizes collagen fibrils and restrains TGF-β signaling, and its downregulation has been linked to disordered matrix assembly and increased fibrotic scarring (Chakravarti, 2002; Zhao, Bai, Xiang, & Pang, 2023; Zheng et al., 2017). In the proteomic dataset, FMOD abundance showed a negative association with age (R² = 0.46, p = 0.0007; Figure 3l). Immunohistochemical (IHC) staining of FMOD in independent ovarian sections from 8 proteomics-matched participants confirmed decreased expression in both cortical and medullary regions of older participants (Figure 3m). Quantification of FMOD IHC further showed a negative correlation with continuous age (R² = 0.62, p = 0.0019; Figure 3n).

### Comparison with ovarian senescence datasets reveals overlap with senescence-associated remodeling

Given the enrichment of inflammatory, extracellular matrix, and secretory programs observed in the native aging ovary, we next asked whether these changes overlapped with ovary-specific senescence-associated molecular signatures. We recognize that acute DNA damage-induced senescence does not fully recapitulate the complexity of chronological ovarian aging. However, we previously showed in an ovarian explant model that doxorubicin-induced senescence signatures showed substantial overlap with molecular features of native ovarian aging (P. R. Devrukhkar et al., 2025). We used independent postmenopausal ovarian senescence datasets to determine whether age-associated proteins identified in this study also overlapped with molecular signatures previously associated with ovarian senescence. These datasets included a senescence-associated secretory phenotype (SASP) proteome from senescence-induced cortex and medulla explants, single-nucleus transcriptomic data from senescence-induced ovary cortex and medulla, and spatial transcriptomic data from p16^INK4a^-positive cortical and medullary regions (P. R. Devrukhkar et al., 2025; Watson et al., 2025). For each comparison, overlap was restricted to proteins or genes meeting the significance thresholds in the corresponding external dataset.

Comparison with the senescence-induced SASP (P. R. Devrukhkar et al., 2025) revealed partial overlap in both ovarian compartments (Figure 4a-b). Between the native proteome and the senescence-induced cortical SASP, 20 proteins overlapped, of which four proteins changed concordantly: Serum amyloid P-component (SAMP), Vitronectin (VTNC), and Alpha-2-macroglobulin (A2MG) were increased, whereas Versican core protein (CSPG2) was decreased (Figure 4a and Figure S6a). A similar pattern was observed with the senescence-induced medulla SASP, where 26 proteins overlapped with the native proteome, and five proteins showed concordant changes: SAMP, VTNC, Protein-glutamine gamma-glutamyltransferase 2 (TGM2), and Nicotinamide phosphoribosyltransferase (NAMPT) were increased, whereas CSPG2 was decreased (Figure 4b and Figure S6b).

**Figure 4.**
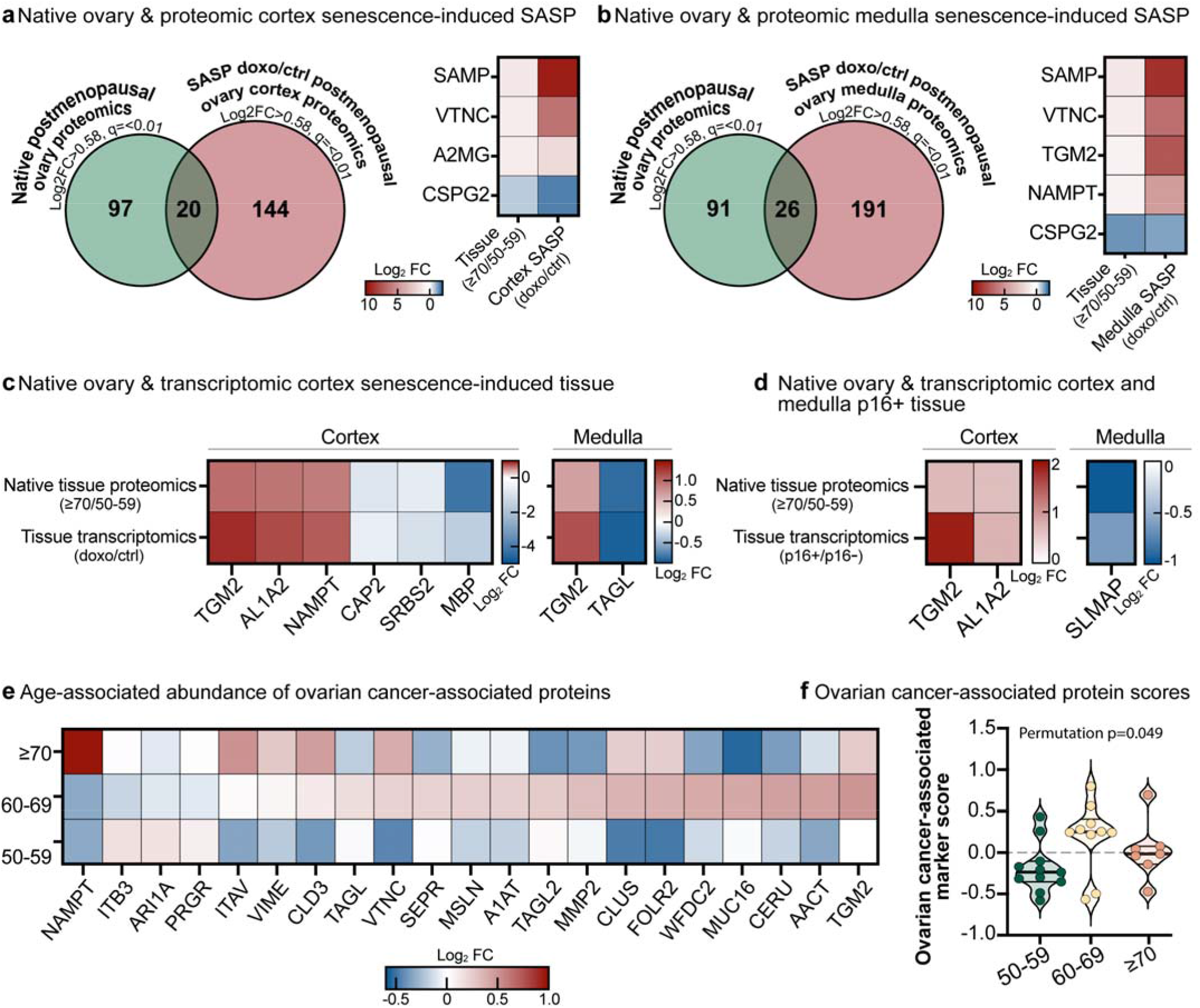
Shared molecular features of postmenopausal ovarian aging, ovarian senescence, and ovarian cancer-associated protein signatures. To place the age-associated proteomic changes identified in the present study into a broader biological context, we compared them with independent postmenopausal ovarian senescence datasets and a literature-curated panel of ovarian cancer-associated proteins. Senescence-induced (Sen-induced) with doxorubicin (doxo), p16+; p16^INK4a^-expressing regions. (**a**) Venn diagram showing overlap between age-associated proteins in native postmenopausal ovary tissue from this study (≥70 vs 50-59-year-old ovaries; log2FC >0.58, q <0.01) and SASP-associated proteomics from postmenopausal ovarian cortex explants treated with doxorubicin (doxo vs control; log2FC >0.5, q <0.05). Of the 20 overlapping proteins, the four showing concordant directional change are displayed in a heatmap. (**b**) Venn diagram showing overlap between age-associated proteins in native postmenopausal ovary tissue from this study (≥70 vs 50-59-year-old ovaries; log2FC >0.58, q <0.01) and SASP-associated proteomics from postmenopausal ovarian medulla explants treated with doxorubicin (doxo vs control; log2FC >0.5, q <0.05). Of the 26 overlapping proteins, the five showing concordant directional change are displayed in a heatmap. (**c**) Overlap between age-associated proteins from this study (≥70 vs 50-59-year-old ovaries; log2FC >0.58, q <0.01) and transcriptomic changes in cortical and medullary explant tissue treated with doxorubicin (doxo vs control; log2FC >0.5, q <0.05). Heatmaps display concordant overlaps (cortex: 6/7; medulla: 2/3). (**d**) Overlap between age-associated proteins from this study (≥70 vs 50-59-year-old ovaries; log2FC >0.58, q <0.01) and transcriptomic differences in cortical and medullary ovarian tissue sections stratified by p16 expression (p16+ vs p16−; log2FC >0.5, q <0.05). Heatmaps display concordant overlaps (cortex: 2/7; medulla: 1/10). (**e**) Heatmap showing cohort mean Log₂ fold abundance values of 21 ovarian cancer-associated proteins identified in the native postmenopausal ovary proteome. Proteins were selected from a literature-curated panel of 35 ovarian cancer-associated proteins, of which 21 were quantified in the present dataset. (**f**) Participant-level ovarian cancer-associated protein scores across age cohorts. Scores were calculated as the mean Log_2_ fold abundance of the 21 quantified ovarian cancer-associated proteins for each participant. Violin plots show the distribution of protein scores for each age group (50-59, 60-69, and ≥70 years). Statistical significance was assessed using a planned permutation test comparing the 60-69-year-old cohort with the combined non-60-69-year-old participants (10,000 permutations).

The native proteome also showed overlap with senescence-induced transcriptomic signatures (P. R. Devrukhkar et al., 2025) (Figure 4c-d). Comparison of the native proteome with the senescence-induced cortical tissue transcriptome identified seven overlapping proteins, six of which changed concordantly: TGM2, Retinal dehydrogenase 2 (AL1A2), and Nicotinamide phosphoribosyltransferase (NAMPT) were increased, whereas Adenylyl cyclase-associated protein 2 (CAP2), Sorbin and SH3 domain-containing protein 2 (SORBS2), and Myelin basic protein (MBP) were decreased (Figure 4c and Figure S6c). Whereas overlap between the native ovarian proteome and the senescence-induced medullary transcriptome was smaller, with three shared proteins and two concordant changes: TGM2 was increased, and Transgelin (TAGLN) was decreased (Figure 4c and Figure S6c). Spatial transcriptomic data from p16^INK4a^-positive regions (Watson et al., 2025) showed a similar trend (Figure 4d). Seven proteins overlapped with the cortical p16^INK4a^-positive regions dataset, of which TGM2 and AL1A2 were concordantly increased, whereas ten proteins overlapped with the medullary p16^INK4a^-positive regions dataset, in which Sarcolemmal membrane-associated protein (SLMAP) was concordantly decreased (Figure 4d and Figure S6d). Together, these comparisons identify a core overlap between native ovarian aging and ovarian senescence-model datasets, characterized by increased SAMP, TGM2, AL1A2, and NAMPT and decreased CSPG2, consistent with a senescence-adjacent remodeling state marked by extracellular matrix remodeling, metabolic stress, and innate immune signaling.

### Healthy ovarian aging exhibits enrichment of ovarian cancer-associated proteomic signatures

Given the enrichment of extracellular matrix remodeling, inflammatory signaling, and senescence-associated molecular programs observed in the aging ovary, we next asked whether these changes also overlapped with proteins previously associated with ovarian cancer. Because senescence-associated tissue remodeling and tumor-associated stroma share several conserved biological processes, including extracellular matrix remodeling, inflammatory signaling, and secretory pathway activation, we assembled a literature-curated panel of 35 proteins reported to be expressed in or functionally implicated in ovarian cancer tissue (Data S17). Twenty-one of these proteins were quantified in our proteomic dataset. Age cohort mean Log_2_ fold protein abundance heatmaps of the matched proteins revealed distinct age-associated patterns across the 50-59, 60-69, and ≥70-year groups (Figure 4e; Data S17). Many ovarian cancer-associated proteins exhibited their highest mean abundance in the 60-69 cohort, including several well-established ovarian cancer markers such as TGM2 (Ibrahim et al., 2025; H. Lee et al., 2026; Zaltron et al., 2024), CLU (M. K. Hassan et al., 2011; Watari et al., 2012), MUC16/CA125 (Huo et al., 2021; C. W. Wang et al., 2024), WFDC2/HE4 (Hamed et al., 2013; Watanabe et al., 2025), and MMP2 (Kenny & Lengyel, 2009; Zeng et al., 2020). Notably, this enrichment occurred in the decade surrounding the median age at ovarian cancer diagnosis (∼63 years) (Caruso, Weroha, & Cliby, 2025; Tew & Lichtman, 2008), suggesting that healthy ovarian tissue in this age range shares molecular features with ovarian cancer-associated proteomic signatures. These findings further suggest a permissive niche concept and the tight association between aging and cancer.

To determine whether this pattern was reflected at the participant level, we next calculated an ovarian cancer-associated protein score for each participant, defined as the mean Log_2_ fold abundance across the 21 detected proteins. Participant-level violin plots demonstrated that the 60-69 cohort had the highest mean protein score (0.19), compared with the 50-59 (-0.19) and ≥70 (0.02) cohorts (Figure 4f). Because our primary hypothesis, based on the cohort heatmap (Figure 4e), was enrichment in the 60-69-year-old group, we performed a planned permutation test comparing participants aged 60-69 years with all remaining participants. The 60-69-year-old cohort exhibited a higher mean protein score than the combined non-60-69-year-old group (0.190 vs. -0.106; difference = 0.296; permutation p = 0.049). Together, these findings demonstrate an age-associated enrichment of ovarian cancer-associated proteins in healthy postmenopausal ovaries, with the strongest enrichment observed in women aged 60-69 years. As all samples analyzed were histologically normal, this enrichment should not be interpreted as evidence of malignancy, but rather as an overlap between healthy ovarian aging and molecular programs previously associated with ovarian cancer.

### Development of the Buck Postmenopausal Ovary Molecular Signature (BuckPOMS)

Collectively, the complementary analytical approaches utilized in this study, including pairwise differential abundance analysis, PLS-DA-derived VIP scores, continuous age-associated linear regression, and age-associated matrisome analysis, identified a recurrent set of proteins consistently associated with postmenopausal ovarian aging. To integrate these findings into a concise molecular framework, we developed the Buck Postmenopausal Ovary Molecular Signature (BuckPOMS), a 26-protein consensus molecular signature representing the most robust age-associated proteomic changes observed across the independent analyses (Figure 5a). The 26 proteins comprising BuckPOMS are displayed as their mean log_2_ abundance across the three postmenopausal age cohorts (Figure 5b & Data S18). The left panel contains proteins that decrease with advancing age, including proteins associated with lipid metabolism and transport (MTP, FABP5, GALC), extracellular matrix and structural organization (FMOD, CPXM1, K2C80), and cellular organization or differentiation (DPYL1, MBP, PCD16). In contrast, the right panel contains proteins that increase with age, including several extracellular matrix remodeling and secreted signaling proteins (ATS16, SNED1, H6ST2, WNT4, CCN5, NDP, IBP5, and WFKN2), together with proteins involved in retinoid metabolism (AL1A2), extracellular nucleotide metabolism (ENPP1), and inflammatory or acute-phase responses (SAMP). These coordinated directional changes highlight a shift away from structural and metabolic maintenance toward increased extracellular remodeling and intercellular signaling across the postmenopausal decades. Together, these coordinated changes comprise BuckPOMS and capture the transition from a proteome enriched in structural maintenance and cellular homeostasis toward one characterized by extracellular matrix remodeling, altered intercellular signaling, and inflammatory remodeling during postmenopausal aging (Figure 5d).

**Figure 5.**
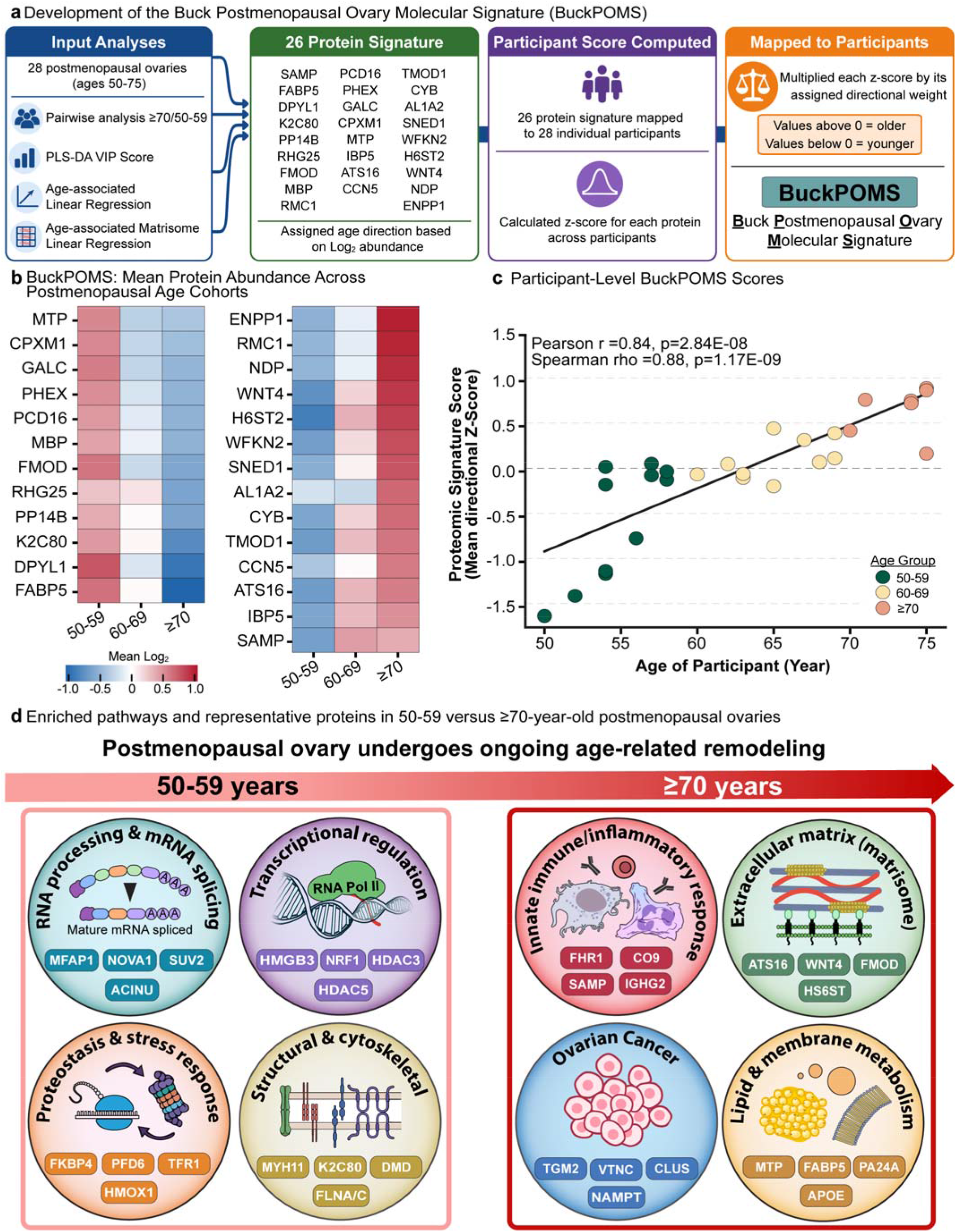
Development and validation of the Buck Postmenopausal Ovary Molecular Signature (BuckPOMS). (**a**) Development of the Buck Postmenopausal Ovary Molecular Signature (BuckPOMS). Candidate proteins were selected by integrating four complementary analytical approaches, including pairwise differential abundance analysis, PLS-DA variable importance in projection (VIP), continuous age-associated linear regression, and age-associated matrisome analyses, to identify the most robust markers of postmenopausal ovarian aging. Twenty-six consensus proteins were retained, assigned an age direction, standardized as Z-scores, and combined into a participant-level BuckPOMS score using directional weighting. (**b**) Heatmap showing the 26 proteins comprising BuckPOMS and their mean log_2_ abundance across postmenopausal age cohorts (50-59, 60-69, and ≥70 years). Proteins in the left panel decrease with advancing age, whereas those in the right panel increase, demonstrating coordinated molecular remodeling across the postmenopausal decades. (**c**) Participant-level BuckPOMS scores calculated as the mean directional Z-score of the 26 signature proteins and plotted against participant age. Positive scores indicate greater similarity to the older postmenopausal proteomic phenotype, whereas negative scores indicate greater similarity to the younger postmenopausal phenotype. Pearson and Spearman correlation coefficients are shown. (**d**) Integrated summary schematic of postmenopausal ovary aging signatures. Pathways and representative proteins enriched in the 50-59-year-old cohort versus the ≥70-year-old cohort are summarized; pathways enriched in one age group are correspondingly depleted in the other (e.g., RNA processing & mRNA splicing is enriched in 50-59-year-olds but reduced in ≥70-year-olds).

To determine whether BuckPOMS captured progressive molecular aging at the individual participant level, protein abundances were standardized as Z-scores, assigned directional weights according to their age-associated expression patterns, and combined to generate a participant-level BuckPOMS score. BuckPOMS scores increased progressively with chronological age (Figure 5c & Data S18), demonstrating a strong positive correlation with age (Pearson r=0.837, P=2.84E-08; Spearman rho=0.875, P=1.17E-09). Consistent with this continuous relationship, BuckPOMS scores also differed significantly across postmenopausal age cohorts (one-way ANOVA, F (2,25)=17.03, P=2.1E-05; η^2^=0.58; permutation P=1.0E-05), indicating that the signature robustly distinguishes younger and older postmenopausal ovaries while capturing progressive molecular remodeling throughout the postmenopausal decades.

Finally, we integrated the major age-associated pathway enrichments identified across our analyses into a summary of postmenopausal ovarian remodeling (Figure 5d). Several of these biological processes were also represented by proteins within BuckPOMS, demonstrating convergence between the consensus signature and the broader pathway-level changes identified across the study. Pathways enriched in ovaries from women aged 50-59 years included RNA processing and mRNA splicing, transcriptional regulation, proteostasis and stress response, and structural and cytoskeletal organization. In contrast, ovaries from women aged ≥70 years were enriched for innate immune and inflammatory responses, extracellular matrix remodeling, ovarian cancer-associated proteins, and lipid and membrane metabolism. Representative proteins associated with each enriched biological process are shown within each category; individual proteins may increase or decrease in abundance irrespective of the overall direction of pathway enrichment. Together, these findings summarize a broader transition from pathways associated with cellular maintenance and structural organization toward immune, extracellular matrix, and metabolic remodeling across the postmenopausal decades.

## Discussion

The postmenopausal ovary has traditionally been viewed as a largely inert fibrotic tissue remnant of reproductive life. Our findings challenge this paradigm. Using proteome-wide data-independent acquisition mass spectrometry using parallel accumulation-serial fragmentation (diaPASEF) to profile non-pathological ovarian tissue across three decades of postmenopausal age, we show that the human ovary continues to undergo substantial molecular remodeling long after reproductive cessation. This work provides significant value and adds complementary knowledge to existing human studies. For example, Ouni et al. (Emna Ouni et al., 2022; E. Ouni et al., 2019) focused on extracellular matrix remodeling from prepubertal life through menopause using an extracellular matrix enrichment strategy. In contrast, our study employed deep quantitative diaPASEF proteomics to profile both extracellular and intracellular proteins, quantifying 5,812 protein groups in histologically normal ovaries collected exclusively from postmenopausal women aged 50-75 years. This design enabled us to specifically investigate whether global molecular remodeling continues across the decades following reproductive cessation. Consistent with Ouni et al., we identified age-associated changes in extracellular matrix and immune-related proteins. However, our broader proteomic coverage further revealed complement activation, inflammatory signaling, cytoskeletal remodeling, and age-associated enrichment of ovarian cancer-associated protein signatures. Together, these findings extend previous studies of the menopausal ovary by demonstrating that molecular remodeling continues throughout later postmenopausal life and support an emerging view of the postmenopausal ovary as a biologically dynamic rather than static tissue.

Across 5,812 quantified proteins, multiple orthogonal analyses converged on a consistent picture of ovarian aging and were integrated into the Buck Postmenopausal Ovary Molecular Signature (BuckPOMS), a 26-protein consensus molecular signature marked by loss of structural and epithelial-associated programs and gain of extracellular matrix remodeling, inflammatory signaling, and complement activation. Ovaries from younger postmenopausal women were enriched for RNA metabolism and mRNA splicing pathways, suggesting a proteome biased toward gene regulation and cellular maintenance, whereas the relative loss of these programs in ovaries from older women was consistent with a shift toward a remodeled, inflammatory tissue state. Importantly, the convergence of independent statistical and biological analyses on the same core set of proteins increases confidence that these changes represent robust features of chronological postmenopausal ovarian aging rather than artifacts of any single analytical approach. One of the clearest signatures of postmenopausal ovarian aging in our dataset was extracellular matrix remodeling. Histological studies have long described fibrotic remodeling in aging ovaries, characterized by stromal expansion and collagen deposition following follicle depletion. Our proteomic data extend these observations by showing that extracellular remodeling continues to evolve across later decades of life, well beyond the initial menopausal transition. In each of the respective analyses, the aging ovary showed a progressive shift toward a matrix-remodeling state enriched for matrisome pathways. This pattern was supported by changes in several extracellular matrix proteins with known structural or stromal roles. Structural matrix components, including fibromodulin (FMOD) and the basement membrane proteoglycan Perlecan (HSPG2), declined with age, whereas remodeling-associated proteins such as transglutaminase-2 (TGM2), A disintegrin and metalloproteinase with thrombospondin motifs 16 (ATS16), vitronectin (VTNC), and Elastin Microfibril Interfacer 3 (EMILIN3), increased. This transition was accompanied by reduced abundance of multiple keratins and other structural proteins, consistent with loss of epithelial and tissue architectural integrity alongside active extracellular remodeling. This pattern is particularly evident in the reciprocal behavior of FMOD and TGM2. FMOD is a small leucine-rich proteoglycan that regulates collagen fibril organization and modulates TGF-β signaling (Chakravarti, 2002; Han et al., 2022; Zhao, Bai, Xiang, & Pang, 2023; Zheng et al., 2017), whereas TGM2 promotes extracellular matrix crosslinking and stabilization during wound repair and fibrosis (Belkin, 2011; Soltani & Kaartinen, 2023; Telci & Griffin, 2006). Together, the decline in FMOD and increase in TGM2 suggest a shift from matrix organization toward a more crosslinked and remodeling-prone stromal environment in older ovaries, a pattern that was further accentuated in our fibrosis analysis within p16 ^INK4a^-positive regions (Watson et al., 2025). A decrease in FMOD has been linked to disrupted matrix assembly and fibrotic remodeling in other tissues, supporting the idea that its loss may contribute to altered stromal architecture in the aging ovary (Ameye et al., 2002; Gill, Oldberg, & Reinholt, 2002; Jiang et al., 2025; E. J. Lee et al., 2021; Zheng, Granado, & Li, 2023). Changes in HSPG2 further indicate disruption of basement membrane architecture, which is notable given perlecan’s role in ovarian epithelial and vascular microenvironments (Hassell et al., 1980; Hayes, Farrugia, Biose, Bix, & Melrose, 2022; Irving-Rodgers & Rodgers, 2005). Increased EMILIN3, a regulator of elastic and microfibrillar matrix organization (Schiavinato et al., 2012; Schiavinato et al., 2024), supports the hypothesis of active stromal reprogramming rather than passive structural deterioration. Taken together, these findings support a model in which the postmenopausal ovary undergoes ongoing extracellular matrix remodeling across later life, consistent with fibrotic and repair-like processes that may continue to reshape ovarian tissue long after reproductive cessation.

In addition to extracellular matrix remodeling, the aging ovarian proteome exhibited features consistent with senescence-adjacent remodeling. Ovaries from older participants were enriched for complement components and inflammatory signaling pathways. These signatures are commonly seen in aging tissues and often described as part of “inflammaging” (Franceschi et al., 2000; Franceschi & Campisi, 2014). Interestingly, our recently published study of post-reproductive ovaries in mice identified a similar post-reproductive molecular signature, characterized by extensive extracellular matrix remodeling and a transition from a reproductive to an immune-like organ state (Converse et al., 2026; Dipali et al., 2023). This cross-species convergence suggests that coordinated extracellular and immune remodeling may represent a conserved feature of the post-reproductive ovary.

Alongside these inflammatory signals, several proteins associated with chromatin regulation and cell cycle control decreased with age. Notable examples include High-Mobility Group Box 3 (HMGB3), Chromodomain Helicase DNA Binding Protein 8 (CHD8), Lysine-specific histone demethylase 1B (KDM1B), Mediator of RNA polymerase II transcription subunit 14 (MED14), and cyclin-dependent kinase 6 (CDK6). These proteins participate in chromatin remodeling, transcriptional regulation, and cell cycle progression, processes that often decline as cells transition toward senescence. HMGB3, a protein that organizes chromatin and is enriched in proliferative cell states, is known to decrease during cellular senescence (Chikhirzhina, Tsimokha, Tomilin, & Polyanichko, 2024; Guo et al., 2016; Nemeth, Kirby, & Bodine, 2006). Meanwhile, CDK6, a cyclin-dependent kinase that promotes G1-S phase cell cycle progression, is suppressed during senescence-associated cell cycle arrest (Crozier et al., 2023; Rajesh et al., 2025; Sherr & Roberts, 1999). Consistent with these observations, comparison with independent ovarian senescence datasets revealed partial overlap between healthy ovarian aging and ovarian senescence-associated molecular signatures (P. R. Devrukhkar et al., 2025; Watson et al., 2025). Proteins, including TGM2, Serum Amyloid P-component (SAMP), Aldehyde dehydrogenase 1 family member A2 (AL1A2), and Nicotinamide phosphoribosyltransferase (NAMPT), were increased, whereas Versican (CSPG2) was decreased with age. These proteins participate in extracellular remodeling, metabolic stress responses, and innate immune signaling, processes characteristic of the senescence-associated secretory phenotype (SASP) (Basisty et al., 2020; Coppe et al., 2008). Several factors linked to secretory programs associated with senescence were identified when comparing ≥70 versus 50-59-year-old ovaries. These included NAMPT, Insulin-like growth factor-binding protein 5 (IGFBP5), TGM2, Clusterin (CLU), Alpha-2-macroglobulin (A2M), and vitronectin. Overall, these observations suggest that the aging postmenopausal ovary acquires a molecular proteomic profile marked by both extracellular remodeling and inflammatory signaling, as well as diminished gene regulatory and growth capacity, with some overlap in programs related to senescence.

The experimentally induced senescence datasets were included to identify molecular features consistently associated with ovarian senescence rather than to serve as surrogates for chronological aging. Although acute DNA damage-induced senescence does not fully recapitulate the complexity of natural ovarian aging, the convergence of these datasets with independent native p16^INK4a^-positive ovarian tissue and our age-associated proteomic changes supports the existence of shared senescence-associated remodeling programs in the postmenopausal ovary.

A recurring theme emerging from BuckPOMS and the complementary analytical approaches was that age-associated extracellular matrix remodeling and complement-innate immune activation led to increasing overlap between the healthy postmenopausal ovary and molecular signatures associated with ovarian cancer. Supporting this observation, we identified enrichment of a literature-curated ovarian cancer-associated protein signature, with the strongest enrichment occurring in women aged 60-69-years-old, near the median age of ovarian cancer diagnosis. Importantly, because all tissues examined were histologically normal, this overlap should not be interpreted as evidence of malignancy, but rather as shared biological processes between healthy ovarian aging and ovarian cancer-associated tissue remodeling.

Consistent with this broader signature, proteins traditionally associated with invasion, chemoresistance, and metastasis in ovarian and other solid cancers, including TGM2 (Ibrahim et al., 2025; H. Lee et al., 2026; Zaltron et al., 2024), VTN (Carreiras, Cruet, Staedel, Sichel, & Gauduchon, 1999; Kenny, Kaur, Coussens, & Lengyel, 2008; Schneider, Suszynska, Kakar, & Ratajczak, 2016), CLU (M. K. Hassan et al., 2011; Watari et al., 2012), and NAMPT (Kudo et al., 2025; Kudo et al., 2020; Sampath, Zabka, Misner, O’Brien, & Dragovich, 2015; Shackelford, Bui, Coppola, & Hakam, 2010), increased with age in the postmenopausal ovary. These findings align with the established roles of these proteins in extracellular matrix remodeling, epithelial-mesenchymal transition (EMT), angiogenesis, and platinum resistance in epithelial ovarian cancer. This change was reinforced by increased matrisome and SASP-like factors such as Asporin (ASPN), A2M, SAMP, IGFBP5, Wnt family member 4 (WNT4), and Heparan-sulfate 6-O-sulfotransferase 2 (H6ST2). These factors indicate cancer-associated fibroblast (CAF)-like ECM stiffening, growth-factor presentation, and metastatic behaviour. HS6ST2 has been identified in pan-cancer analyses as a glycosaminoglycan-modifying enzyme (Chen et al., 2023; Huang et al., 2024). HS6ST2 overexpression is linked to more advanced disease stages and poorer overall survival across multiple tumor types, highlighting its potential as a prognostic marker in cancer (Backen et al., 2007; Chen et al., 2023; Cole, Rushton, Jayson, & Avizienyte, 2014; Huang et al., 2024).

In parallel, complement and innate immune pathways increased with age. Previous research shows that ECM and inflammatory signals in the peritoneal-ovarian niche actively promote attachment, invasion, and macrophage recruitment rather than acting as a passive scaffold (Ricciardelli, Lokman, Ween, & Oehler, 2016). In this context, the loss of structural smooth-muscle and keratin networks, along with a decline in matrix-organizing proteins such as Fibromodulin (FMOD), likely further disrupts normal tissue architecture. This enables malignant cells from the fallopian tube or peritoneum to implant and grow. Our data support the view that, even in the absence of malignancy, the aging postmenopausal ovary moves toward a more “cancer-ready” niche. This adds complexity to the discussion around prophylactic removal of ovaries (salpingo-oophorectomy). While risk-reducing surgery clearly lowers ovarian cancer incidence and mortality in high-risk BRCA1/2 carriers (Liu et al., 2022; Rebbeck, Kauff, & Domchek, 2009), early or indiscriminate removal of ovaries comes with higher long-term risks of cardiovascular disease, osteoporosis, and cognitive decline, especially if performed close to the natural age of menopause (H. Hassan et al., 2024; Parker, Jacoby, Shoupe, & Rocca, 2009; Rivera et al., 2009).

Beyond its potential implications for ovarian cancer susceptibility, our findings suggest that the postmenopausal ovary follows remodeling trajectories similar to those observed across multiple aging tissues. These include loss of structural integrity, extracellular matrix remodeling, activation of innate immune pathways, and metabolic adaptation. However, compared with organs such as muscle, liver, brain, and immune tissues, the aging ovary has remained relatively understudied. Our proteome-scale analysis begins to position the ovary within the broader landscape of human tissue aging.

Overall, our findings suggest that decisions regarding ovary removal in older women should consider not only the established cancer-preventive benefits of oophorectomy but also the possibility that removal of a postmenopausal ovary undergoing persistent fibroinflammatory remodeling may have broader implications for systemic aging and age-related disease. However, because the present study was not designed to evaluate clinical outcomes, whether these age-associated molecular changes translate into meaningful health risks remains an important future direction. Instead, the molecular pathways identified here provide a framework for further research to investigate how ongoing remodeling of the postmenopausal ovary may influence systemic aging and susceptibility to age-related diseases in women. By focusing on healthy, histologically normal postmenopausal ovaries across successive decades of life, this study establishes a reference map of chronological ovarian aging that can serve as a foundation for future investigations into disease-associated remodeling.

A limitation of this study is that we did not capture age at natural menopause; years since menopause, and menopausal hormone therapy history were unavailable for this cohort. Ovarian histology independently confirmed a postmenopausal state, including the absence of follicles; these additional clinical variables may influence ovarian remodeling after reproductive cessation. Moreover, accurately determining menopausal timing is challenging because menopausal status and age at menopause are frequently based on self-report, while surgical history and menopausal hormone therapy can further complicate classification(de Vries, Northington, Kaye, & Bogner, 2011; den Tonkelaar, 1997; Pieterse, Picavet, MenoPause, Verschuren, & Broekman, 2026; Yap et al., 2022). Accordingly, our analyses characterize molecular remodeling across the postmenopausal decades rather than the menopausal transition itself or the effects of exogenous hormone exposure. Despite these limitations, our findings demonstrate that chronological aging after menopause is accompanied by progressive molecular remodeling of the human ovary, supporting the concept that the postmenopausal ovary remains a biologically active organ rather than a static residual tissue. Future prospective studies incorporating accurately documented menopausal timing, hormone therapy history, and longitudinal follow-up will be important for distinguishing ovarian changes associated with the menopausal transition, time since menopause, and chronological aging after reproductive cessation.

These findings challenge the paradigm that the postmenopausal ovary is biologically inert. Instead, the ovary appears to be an actively remodeling tissue, characterized by changes in the extracellular matrix, signals related to inflammation, and altered metabolic programs. The age-associated increase in several secreted and extracellular factors, including IGFBP5, NAMPT, CLU, vitronectin, and WNT4, raises the possibility that the postmenopausal ovary continues to contribute to local tissue signaling and potentially to broader organ-organ communication during aging. Overall, our data, summarized in the 26-protein Buck Postmenopausal Ovary Molecular Signature (BuckPOMS), support a model in which the postmenopausal ovary is not a static fibrotic remnant of reproductive life, but an actively remodeling tissue with the potential to participate in inter-organ signaling during aging. As a consensus molecular signature derived from multiple complementary analytical approaches, BuckPOMS provides a proteomic framework for future studies investigating the mechanisms, biomarkers, and clinical consequences of postmenopausal ovarian aging, senescence, and age-associated ovarian disease.

## Materials and Methods

### Human Ovarian Tissue Acquisition and Processing

De-identified human ovarian tissue was obtained from the Northwestern University Reproductive Tissue Library (NU-RTL) under Institutional Review Board-approved protocols (STU00215770). Ovaries were obtained from females aged 50-75Oyears old (Average 63.2O±O7.9Oyears) undergoing bilateral salpingo-oophorectomies and/or total laparoscopic hysterectomies for various gynecologic conditions with no malignant pathology (Data S1). Upon collection, the tissue was divided into cross-sections (3-5Omm thick) that were generated perpendicular to the long axis of the ovary. In the absence of significant gross pathology as assessed by a certified gynecologic pathologist, up to two ovarian cross-sections were designated for research and transported to the laboratory on ice in ORIGIO Handling IVF medium (Cooper Surgical Inc., Trumbull, CT, USA) (P. A. Devrukhkar, H.; Giornazi, A.; Shah, S.; Pavone, M.E.; Duncan, F.E. , 2023). For proteomics analysis, ovarian tissue samples (20-30 mg) were flash-frozen and stored at -80°C until processing. For histological analysis, ovarian tissue pieces were fixed in Modified Davidson’s Fixative (Electron Microscopy Sciences, Hatfield, PA, 64133-50) for 2 hours at room temperature and then overnight at 4°C. Tissue pieces were washed in 70% ethanol, transferred to 70% ethanol, and stored at 4°C until further processing. Tissue pieces were then dehydrated in an automated tissue processor (Leica Biosystems, Wetzlar, Germany, TP1020), embedded in paraffin, and sectioned (5 µm thickness) with a microtome (Leica Biosystems, RM2155).

### Tissue Lysis and Homogenization

Human postmenopausal ovarian tissue from 28 participants ranging from 50-75 years of age was prepared for proteomic analysis. Samples were homogenized in ∼300 µL lysis buffer containing 8 M urea, 2% sodium dodecyl sulfate (SDS), 1 µM trichostatin A (TSA), 3 mM nicotinamide adenine dinucleotide (NAD), 75 mM sodium chloride, and 1X protease and phosphatase inhibitor cocktail (Thermo Fisher Scientific, Waltham, MA) in 200 mM triethylammonium bicarbonate (TEAB) by adding them to 2.0 mL safe-lock tubes (VWR International, Radnor, PA) containing stainless steel beads and subjected to three intervals of high-speed shaking (25 Hz, 1 min) using a Qiagen TissueLyser II (Qiagen, Hilden, Germany). Tissue homogenates were centrifuged at 15,700 x g for 10 min at 4°C, and the supernatant was collected for label-free quantitative proteomics experiments. Protein concentration was determined using Bicinchoninic Acid (BCA) assay (Thermo Fisher Scientific, Waltham, MA).

### Protein Digestion and Desalting

Aliquots of 100 µg protein lysates for each sample were brought to the same overall volume of 50 µL with water, reduced using 20 mM dithiothreitol in 50 mM TEAB at 50°C for 10 min, cooled to room temperature (RT) and held at RT for 10 min, and alkylated using 40 mM iodoacetamide in 50 mM TEAB at RT in the dark for 30 min. Samples were acidified with 12% phosphoric acid to obtain a final concentration of 1.2% phosphoric acid. S-Trap buffer, consisting of 90% methanol in 100 mM TEAB at pH ∼7.1, was added, and samples were loaded onto the S-Trap micro spin columns. The entire sample volume was spun through the S-Trap micro spin columns at 4,000 x g and RT, binding the proteins to the micro-spin columns. Subsequently, S-Trap micro spin columns were washed twice with S-Trap buffer at 4,000 x g and RT and placed into clean elution tubes. Samples were incubated for one hour at 47 °C with sequencing-grade trypsin (Promega, San Luis Obispo, CA) dissolved in 50 mM TEAB at a 1:25 (w/w) enzyme: protein ratio. An additional aliquot of trypsin dissolved in 50 mM TEAB was added, and samples were digested overnight at 37 °C. Peptides were sequentially eluted from micro-S-Trap spin columns with 50 mM TEAB, 0.5% formic acid (FA) in water, and 50% acetonitrile (ACN) in 0.5% FA. After centrifugal evaporation, samples were resuspended in 0.2% FA in water and desalted with Oasis 10-mg Sorbent Cartridges (Waters, Milford, MA). The desalted elutions were subjected to centrifugal evaporation and re-suspended in 0.1% FA in water at a final concentration of 1 µg/µL.

### Mass Spectrometric Analysis

400 ng of each sample were loaded onto EvoTips Pure (Evosep, Odense, Denmark) following the manufacturer’s protocol. During sample loading onto EvoTips Pure, one microliter of indexed Retention Time Standard (iRT, Biognosys, Schlieren, Switzerland) was added to each sample. HPLC-MS/MS analyses were performed on an Evosep One liquid chromatography system (Evosep) coupled to a timsTOF HT mass spectrometer (Bruker, Bremen, Germany). The solvent system consisted of 0.1% FA in water (solvent A) and 0.1% FA in ACN (solvent B). Peptides were eluted on a PepSep C_18_ analytical column (150 µm x 15 cm, 1.5 µm particle size; Bruker) using the 30 SPD method (44-minute gradient length, 500-nL/min flow rate). A zero-dead volume emitter was installed in the nano-electrospray source (CaptiveSpray source, Bruker Daltonics), and the source parameters were set as follows: capillary voltage 1600 V, dry gas 3 L/min, and dry temperature 180°C. Each sample was acquired in data-independent acquisition (DIA) mode coupled to parallel accumulation serial fragmentation (PASEF) or dia-PASEF mode (Meier et al., 2020) with 1 survey TIMS MS scan and 24 dia-PASEF MS/MS scans per 2.03-s cycle. The dual TIMS analyzer was operated at a 100% duty cycle, with equal accumulation and ramp times of 75 ms each. For each TIMS MS scan, the ion mobility ranged from 1/K_0_ = 0.85-1.30 Vs/cm^2^, and the mass range covered *m/z* 100-1,700. The dia-PASEF scheme was defined as 63 x 15 Th isolation windows from *m/z* 305.9 to 1,250.9 and covering the ion mobility range 1/K_0_ = 0.74-1.30 Vs/cm^2^ (Data S19). The collision energy was defined as a linear function of mobility, starting from 20 eV at 1/K_0_ = 0.6 Vs/cm^2^ to 59 eV at 1/K_0_ = 1.6 Vs/cm^2^. For calibration of the ion mobility dimension, three ions of Agilent ESI-Low Tuning Mix ions were selected (m/z [Th], 1/K0 [Th]: 622.0289, 0.9915; 922.0097, 1.1986; 1221.9906, 1.13934).

### DIA Data Processing and Statistical Analysis

DIA data were processed in Spectronaut v17 (version 17.6.230428.55965) using directDIA (Bruderer et al., 2017; Collins et al., 2017; Gillet et al., 2012). Data extraction parameters were set as dynamic and non-linear iRT calibration, with precision iRT selected (Escher et al., 2012). Data were searched against the *Homo sapiens* reference proteome with 20,380 entries (UniProtKB-TrEMBL), accessed on 01/29/2021. Trypsin/P was set as the digestion enzyme, and two missed cleavages were allowed. Cysteine carbamidomethylation was set as a fixed modification, while methionine oxidation and protein N-terminus acetylation were set as dynamic modifications. Identification was performed using 1% precursor and protein q-value. Quantification was based on the peak areas of extracted ion chromatograms (XICs) of 3-6 MS2 fragment ions, specifically b- and y-ions, with local normalization and q-value sparse data filtering applied. In addition, iRT profiling was selected. Differential protein expression analysis comparing 1) 60-69 vs. 50-59-year-old age groups, and 2) >70 vs. 50-59-year-old age groups was performed using an unpaired t-test, and p-values were corrected for multiple testing using the Storey method (Burger, 2018). Specifically, group-wise testing corrections were applied to obtain q-values. Protein groups with at least two unique peptides, q-value < 0.01, and absolute Log_2_ fold-change > 0.58 are significantly altered (Data S2-5).

### Immunohistochemistry

Slides were deparaffinized in Citrisolv clearing agent (Decon Labs, 1601), rehydrated in decreasing concentrations of ethanol (Fisher Scientific, BP2818-4), and finally rinsed in distilled water. Heat-induced antigen retrieval was performed in citrate buffer (pH 6). Slides were washed two times for 15 min each in Tris-buffered saline (TBS) containing 0.1% Tween-20 (Sigma, P2287) (TBS-T), then incubated in 3% H_2_O_2_ for 15 min to block endogenous peroxidase activity. Slides were rinsed in TBS and blocked using an avidin/biotin blocking kit according to the manufacturer’s instructions (Vector Laboratories, SP-2001). Slides were then rinsed in TBS and incubated in protein block solution (10% serum from the host species of the secondary antibody, 3% Bovine Serum Albumin - Fraction V) in 1X TBS for 45 min at room temperature. The protein block solution was removed, and slides were incubated in the respective primary antibodies diluted in protein block solution overnight in a humidified chamber at 4°C. Primary antibodies used were rabbit anti-FMOD (Thermo Fisher Scientific, PA5-26250; 1:200), and rabbit anti-WNT4 (Proteintech, 14371-1-AP; 1:200). For nonimmune controls, instead of the primary antibody, sections were incubated with species-matched IgG (Rabbit IgG) at the same concentrations as the corresponding primary antibody. Secondary antibody controls were performed by incubating slides with antibody diluent only to evaluate nonspecific binding. Slides were washed three times in TBS-T for 5 min each and then incubated with secondary antibodies for 1 hour at room temperature (Vector Laboratory, PK-6101). Slides were washed three times in TBS-T for 5 min each and then incubated with avidin/biotin complex (ABC) (Vector Laboratory, PK-6101) according to the manufacturer’s instructions. Following the final wash in TBS-T, detection was performed using a DAB Peroxidase (HRP) Substrate Kit (Vector Laboratories, SK-4100) according to the manufacturer’s instructions. DAB substrate reaction was stopped by incubating in distilled water for 5 min. Slides were counterstained with Harris Hematoxylin (Fisher Healthcare,245651) and mounted on glass slides. All tissue sections were processed at the same time using the same immunostaining protocol. Slides were visualized using the ZEISS Axioscan 7 Imaging system with a 20X objective. Image analysis was performed in QuPath (Bankhead et al., 2017). Background subtraction and color deconvolution were applied to separate hematoxylin and DAB signals. A pixel classifier was used to segment total tissue and to identify DAB-positive regions based on color and intensity features. The DAB-positive area was quantified and reported as the percentage of DAB-positive area relative to the total tissue area.

### Post hoc analysis

#### Differential abundance testing and volcano plot visualization

For each pairwise age comparison, differential abundance results were imported from tabular outputs containing average log2 fold change (AVG Log2 Ratio), q-values, and protein identifiers. Volcano plots were generated in Python using standard data manipulation and visualization workflows. The x-axis represents AVG Log2 Ratio, and the y-axis represents −log10(q-value). Proteins were classified as upregulated (log2FC >0.58 and q <0.01), downregulated (log2FC <-0.58 and q <0.01), or not significant (all others). Thresholds were indicated with vertical dashed lines at log2FC = ±0.58 and a horizontal dashed line at q = 0.01. Annotated plots were created by labeling significant proteins using the ProteinNames field.

#### Proteomic preprocessing for multivariate modelling

For multivariate analyses, protein intensities were exported as a pivot table of protein groups, retaining only protein groups supported by ≥2 unique peptides. Protein and gene symbols were used as feature identifiers. Intensities were transformed as log2(intensity + 1) to stabilize variance and reduce the influence of extreme values. Missing values were imputed within each protein using the median intensity of that protein across participants. The final matrix was formatted as samples (rows) x proteins (columns), and each protein was standardized to z-scores (mean 0, standard deviation 1) before modeling.

#### Partial least squares-discriminant analysis (PLS-DA)

To test whether the global ovarian proteome separates by postmenopausal age group, we performed partial least squares-discriminant analysis (PLS-DA) using a two-component PLS regression framework. Only age-group labels (50-59, 60-69, ≥70) were used for classification (i.e., models were not covariate-adjusted). We evaluated a three-class model (all groups) and pairwise contrasts (50-59 vs 60-69; 50-59 vs ≥70). Sample separation was visualized using score plots of the first two latent variables (LV1 and LV2), and group dispersion was summarized with covariance-based ellipses.

#### Cross-validation and ROC analysis for PLS-DA models

Model performance was evaluated using stratified 5-fold cross-validation to preserve class proportions in each split. Within each training fold, features were standardized, and a two-component PLS model was fit, then applied to held-out samples to generate out-of-fold predictions. For the three-class model, accuracy was calculated using the predicted class with the largest model response. For binary comparisons, classification performance was summarized using both accuracy and receiver operating characteristic (ROC) curves computed from pooled out-of-fold predictions; the older age group was treated as the positive class, and the corresponding predicted response was used as a continuous decision variable. The area under the ROC curve (AUC) was reported as a summary measure of discrimination.

#### VIP-based feature prioritization and VIP-LV1 plots

To identify proteins most responsible for age-group separation, we computed variable importance in projection (VIP) scores from the fitted PLS models (Akarachantachote et al., 2014; Chong & Jun, 2005). VIP scores summarize the contribution of each protein to explaining variation in the group label across the modeled components. For visualization, we plotted VIP score against LV1 loading (the primary discriminant axis). Proteins with VIP ≤1.5 were considered non-prioritized, while proteins with VIP >1.5 were highlighted and colored by LV1 direction (positive indicating relatively higher abundance in the older group; negative indicating relatively higher abundance in the younger group). For the 50-59 vs ≥70 comparison, we defined representative VIP-selected proteins as the top 20 older-associated (LV1 >0) and top 20 younger-associated (LV1 <0) proteins ranked by VIP.

#### Heatmap visualization of VIP-selected proteins

Heatmaps were generated to visualize coordinated age-associated shifts among VIP-selected proteins. Where multiple protein groups mapped to the same gene symbol, intensities were aggregated at the gene level using the median. For participant-level heatmaps, log2-transformed intensities were z-scored for each protein across participants, and participants were ordered by chronological age. To summarize group-level patterns, participant-level z-scores were averaged within each age category (50-59, 60-69, ≥70).

#### Covariate-adjusted regression modeling and linear-quadratic comparison

To identify proteins associated with age while accounting for relevant participant characteristics, we performed per-protein ordinary least squares regression with age, BMI, and self-reported race/ethnicity (categorical) as covariates. Proteins were required to have ≥6 non-missing participant values and a nonzero variance. Two nested models were evaluated: Linear age model: abundance ∼ Age + BMI + Race/ethnicity, and Quadratic age model: abundance ∼ Age + Age² + BMI + Race/ethnicity. Model fit was compared using the Akaike Information Criterion (AIC), the Bayesian Information Criterion (BIC), and an F-test for nested model comparison (quadratic vs. linear). For downstream age-association analyses, the preferred model (linear age term) was used. Multiple testing was addressed using Storey q-values (false discovery rate control) for key terms. Log transformation note: For selected analyses and figures, abundances were transformed using log1p, defined as ln(1 + abundance).

#### Gene set enrichment analysis (GSEA)

A pre-ranked gene set enrichment analysis was used to interpret proteomic signatures at the pathway level. For PLS-DA-based enrichment (50-59 vs ≥70), proteins were filtered to VIP >1.5 and ranked using a signed score based on LV1 directionality so that positive ranks corresponded to proteins relatively higher in ≥70 and negative ranks corresponded to proteins relatively higher in 50-59. Pathway enrichment was evaluated using Reactome (Matthews et al., 2009) and Hallmark (Liberzon et al., 2015) gene sets from <u>MSigDB</u>. Pre-ranked GSEA was implemented using gseapy with 1,000 permutations, standard gene set size filters (minimum 10; maximum 2,000), and a fixed random seed. Results are reported as normalized enrichment score (NES) and FDR q-value; NES >0 indicates enrichment toward the ≥70-associated end of the ranked list, and NES <0 indicates enrichment toward the 50-59-associated end.

#### Extracellular matrix-focused enrichment and Matrisome analyses

To test whether extracellular matrix-related proteins increase with age, we performed enrichment analyses using MatrisomeDB (Naba et al., 2012; Shao et al., 2023)-derived gene sets. Enrichment results are reported as NES and FDR q-values, and subcategory-level analyses were used to identify which matrisome components drove the overall signal. In addition, targeted covariate-adjusted regression restricted to matrisome features was used to highlight the most age-associated matrisome proteins.

Visual Studio Code and GraphPad Prism were used for visualization and analysis of data.

## Author Contributions

B.S., F.E.D., M.A.W., and S.M. conceived and designed the study (Conceptualization). E.D.S., M.E.G.P., and P.R.D. oversaw the acquisition of postmenopausal ovarian tissue and pathology assessment (Resources, Investigation). B.So. provided biospecimens, prepared histological sections, and performed histological analyses, including WNT4 and FMOD staining (Resources, Methodology, Investigation). M.A.W processed ovarian samples for mass spectrometry acquisition (Investigation). C.D.K performed data-independent acquisition mass spectrometry using parallel accumulation-serial fragmentation (diaPASEF) on ovarian samples (Methodology, Investigation). M.A.W processed the diaPASEF data in Spectronaut v17 and conducted differential expression, multivariate, covariate, and pathway enrichment analyses (Methodology, Data Curation, Formal Analysis). M.A.W. prepared figures and data visualizations (Visualization). M.A.W. wrote the first draft of the manuscript (Writing, Original Draft). M.A.W., B.S., F.E.D., B.So., and P.R.D. critically revised and edited the manuscript (Writing, Review & Editing). M.A.W., B.S., and F.E.D. provided supervision and project oversight (Supervision). All authors read and approved the final manuscript.

## Acknowledgments

We gratefully acknowledge the women who generously donated ovarian tissue for this study. Their contribution made this work possible and advances our understanding of postmenopausal ovarian aging and women’s health. We thank Asma Giornazi, Dr. Tomiris Atazhanova, and Uyen Tran at Northwestern University for their critical roles in tissue acquisition. We acknowledge the commitment and scientific engagement of the late Dr. Judy Campisi and her passionate support of this project.

## Data Availability

Raw data and complete MS data sets have been uploaded to the Mass Spectrometry Interactive Virtual Environment (MassIVE) repository, developed by the Center for Computational Mass Spectrometry at the University of California, San Diego, and can be downloaded (username and password please see below) using the following link: <u>Massive Link</u> (MassIVE ID Number: MSV000101019; ProteomeXchange ID: PXD075113). The processed proteomics data and complete Supplementary Data workbook (Data S1-S19) are publicly available through Zenodo (https://doi.org/10.5281/zenodo.21880448).

## Ethical Statement & Consent

De-identified human ovarian tissue was obtained from the Northwestern University Reproductive Tissue Library (NU-RTL) under Institutional Review Board–approved protocols (STU00215770). Ovaries were collected from women aged 50-75 years (mean 63.2O±O7.9 years) undergoing bilateral salpingo-oophorectomy and/or total laparoscopic hysterectomy for benign gynecologic indications with no malignant pathology (Data S1). All tissue donations were made under written informed consent, and all procedures were conducted in accordance with the Declaration of Helsinki and relevant institutional guidelines and regulations.

## Supplemental Figures

Figure S1. Three-way PLS-DA comparison and WNT4 IHC quantification. (a) Three-group (50-59, 60-69, ≥70-year-old) PLS-DA comparison. Score plots show the first two latent variables (LV1 and LV2); each point represents a participant, and ellipses indicate group dispersion in LV space. (b) Quantification of WNT4 IHC staining across 10 participants was associated with participant age. Linear regression is shown with 95% confidence interval (shaded); R² and p-value are indicated.

Figure S2. Covariate-adjusted model comparison indicates the linear age model outperforms the quadratic age model. Several complementary metrics were used to compare covariate-adjusted linear and quadratic age models across proteins. (a) Histogram of Akaike Information Criterion (AIC) differences, shown as ΔAIC = AIC (quadratic) – AIC (linear) for each protein. Values to the right of 0 indicate a higher AIC (better fit) for the linear model, whereas values to the left favor the quadratic model. Arrows annotate the number of proteins on each side of ΔAIC = 0. (b) Histogram of Bayesian Information Criterion (BIC) differences, shown as ΔBIC = BIC (quadratic) – BIC (linear) for each protein. Values to the right of 0 indicate a higher BIC (better fit) for the linear model, whereas values to the left favor the quadratic model. Arrows annotate the number of proteins on each side of ΔBIC = 0. (c) Summary of model preference, showing the number of proteins favoring the linear versus quadratic model based on AIC and BIC. (d) Scatter plot comparing ΔAIC with the significance of the quadratic term assessed by an F-test (−log10 p-value). Negative ΔAIC values favor the quadratic model. The horizontal line denotes p = 0.05. Proteins in the upper-left quadrant (ΔAIC <0 and p <0.05) are those for which the quadratic model is supported by both improved information criterion and a significant F-test. Numbers within each quadrant indicate protein counts.

Figure S3. Top age-associated proteins from covariate-adjusted linear regression. (a) Forest plot showing β_age estimates (±95% CI) for the two significant age-associated proteins (q <0.05) along with the 28 top-ranked candidate proteins (q <0.4). (b) Forest plot showing the corresponding percent change in protein abundance per decade, derived from β_age, for the two significant proteins (q <0.05) along with the 28 top-ranked candidate proteins (q <0.4).

Figure S4. Pathway clustering of top proteins from the covariate-adjusted linear regression model. The top 30 proteins from the covariate-adjusted linear regression model, ranked by q-value, are grouped into functional pathway clusters (2 significant proteins, q <0.05; 28 additional top-ranked candidates, q < 0.4). For each protein, ln(1 + abundance) is plotted against age across participants, with the fitted linear regression line and 95% confidence interval (shaded); R² and p-value are shown. Proteins are organized into the following pathway clusters: (a) Extracellular matrix organization & remodeling. (b) Lipid metabolism & secretory trafficking. (c) Gene regulation & cell cycle control. (d) Cytoskeletal dynamics & membrane trafficking.

Figure S5. Covariate-adjusted analysis reveals age-associated pathways in the postmenopausal ovaries. (a) A bubble plot summarizing gene set enrichment analysis (GSEA) using MSigDB Reactome and Hallmark gene sets, performed on age association scores derived from the covariate-adjusted linear model fit to ln (1 + abundance). Pathways are ranked by q-value; the top 4 pathways meet q <0.05. (b-d) These panels highlight representative pathways prioritized from the global enrichment results in (a), reporting normalized enrichment score (NES) and FDR (q-value) to assess age-associated enrichment: (b) keratinization, (c) hedgehog signaling, and (d) inflammatory response.

Figure S6. Cross-validation with ovarian senescence datasets. To assess concordance with prior postmenopausal ovarian senescence datasets, we tested whether age-associated proteins identified in the current study overlapped with senescence-associated proteomic and transcriptomic signatures from independent ovarian explant and tissue-section models. Sen-induced; senescence-induced with doxorubicin (doxo), p16+; p16^INK4a^ expressing regions. (a) Heatmap showing all 20 overlapping proteins between age-associated proteins in native postmenopausal ovary tissue from this study (≥70 vs 50-59-year-old; log2FC >0.58, q <0.01) and SASP-associated proteomics from postmenopausal ovarian cortex explants treated with doxorubicin (doxo vs control; log2FC >0.5, q <0.05). (b) Heatmap showing all 26 overlapping proteins between age-associated proteins in native postmenopausal ovary tissue from this study (≥70 vs 50-59-year-old; log2FC >0.58, q <0.01) and SASP-associated proteomics from postmenopausal ovarian medulla explants treated with doxorubicin (doxo vs control; log2FC >0.5, q <0.05). (c) Venn diagrams showing overlap between age-associated proteins in native postmenopausal ovary tissue from this study (≥70 vs 50-59-year-old; log2FC >0.58, q <0.01) and transcriptomic changes in cortical and medullary explant tissue treated with doxorubicin (doxo vs control; log2FC >0.5, q <0.05). Heatmaps of all overlapping cortical and medullary proteins are shown. (d) Venn diagrams showing overlap between age-associated proteins in native postmenopausal ovary tissue from this study (≥70 vs 50-59-year-old; log2FC >0.58, q <0.01) and transcriptomic differences in cortical and medullary ovarian tissue sections stratified by p16 expression (p16+ vs p16-; log2FC >0.5, q <0.05). Heatmaps of all overlapping cortical and medullary proteins are shown.

