## Supplementary material for "The human ovary exhibits dynamic molecular remodeling in the decades post-menopause": Suppl Figure S1

### Slide 1
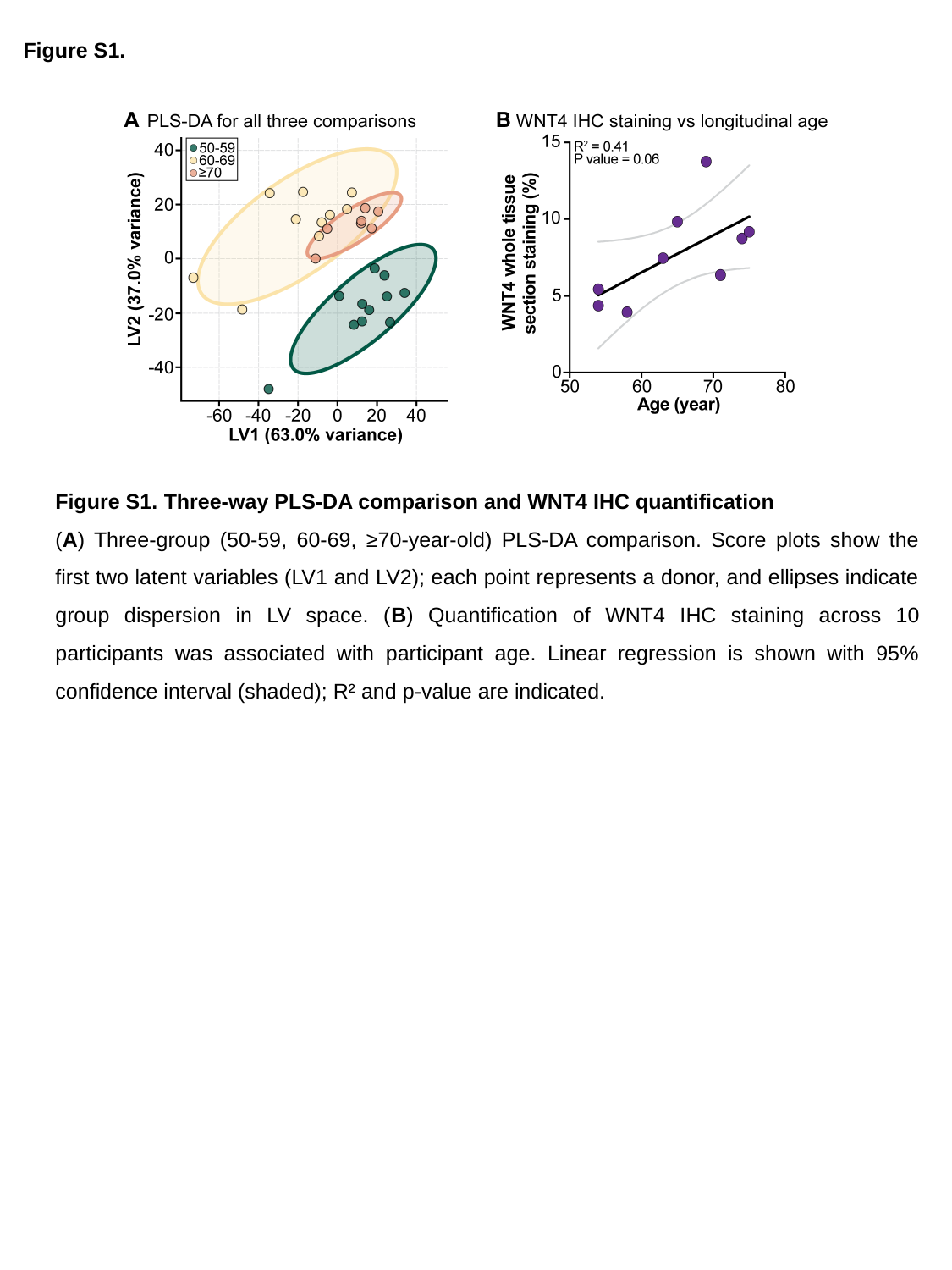

Figure S1.
Figure S1. Three-way PLS-DA comparison and WNT4 IHC quantification
(A) Three-group (50-59, 60-69, ≥70-year-old) PLS-DA comparison. Score plots show the first two latent variables (LV1 and LV2); each point represents a donor, and ellipses indicate group dispersion in LV space. (B) Quantification of WNT4 IHC staining across 10 participants was associated with participant age. Linear regression is shown with 95% confidence interval (shaded); R² and p-value are indicated.
