## Supplementary material for "The human ovary exhibits dynamic molecular remodeling in the decades post-menopause": Suppl Figure S2

### Slide 1
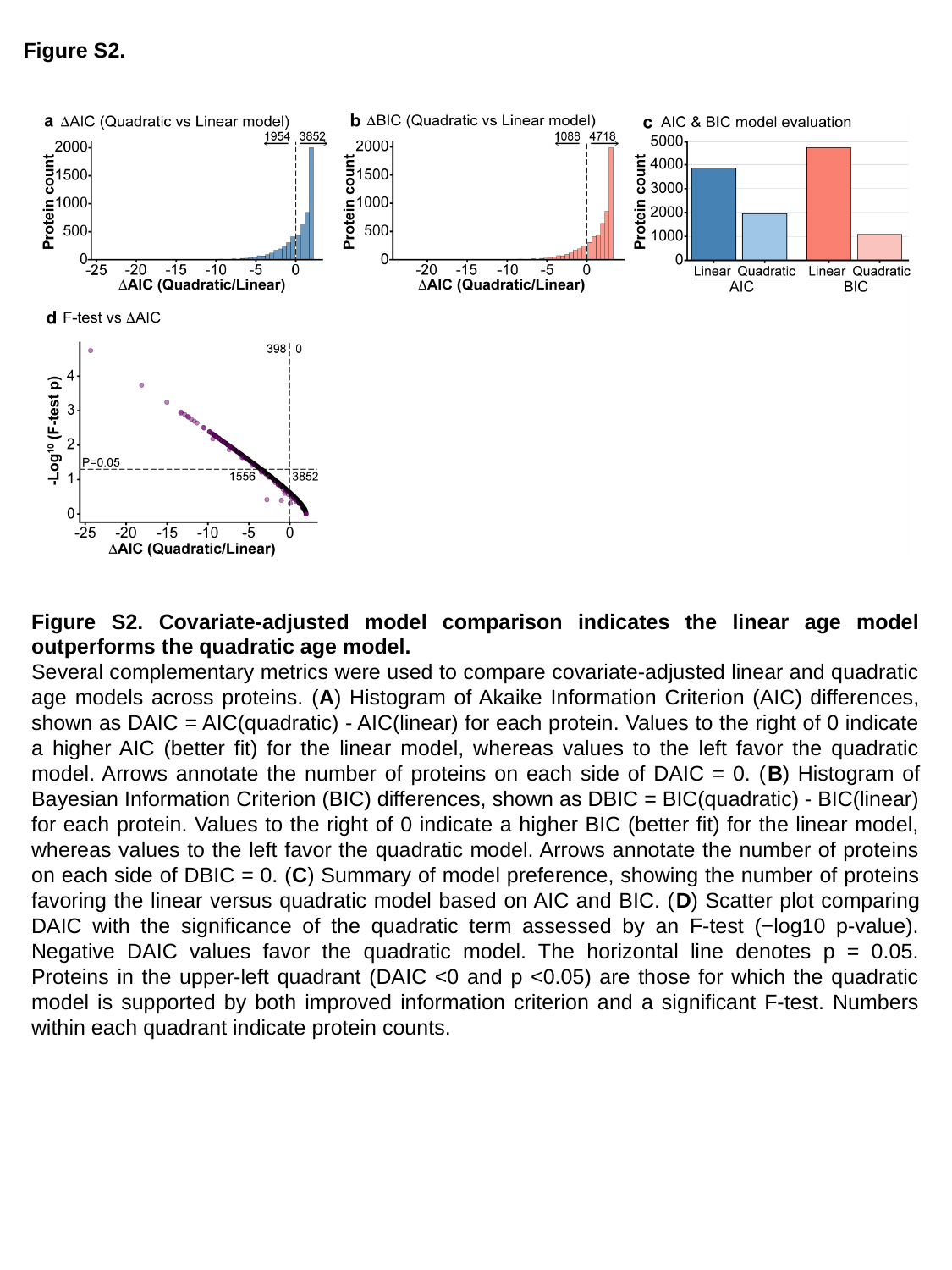

Figure S2.
Figure S2. Covariate-adjusted model comparison indicates the linear age model outperforms the quadratic age model.
Several complementary metrics were used to compare covariate-adjusted linear and quadratic age models across proteins. (A) Histogram of Akaike Information Criterion (AIC) differences, shown as DAIC = AIC(quadratic) - AIC(linear) for each protein. Values to the right of 0 indicate a higher AIC (better fit) for the linear model, whereas values to the left favor the quadratic model. Arrows annotate the number of proteins on each side of DAIC = 0. (B) Histogram of Bayesian Information Criterion (BIC) differences, shown as DBIC = BIC(quadratic) - BIC(linear) for each protein. Values to the right of 0 indicate a higher BIC (better fit) for the linear model, whereas values to the left favor the quadratic model. Arrows annotate the number of proteins on each side of DBIC = 0. (C) Summary of model preference, showing the number of proteins favoring the linear versus quadratic model based on AIC and BIC. (D) Scatter plot comparing DAIC with the significance of the quadratic term assessed by an F-test (−log10 p-value). Negative DAIC values favor the quadratic model. The horizontal line denotes p = 0.05. Proteins in the upper-left quadrant (DAIC <0 and p <0.05) are those for which the quadratic model is supported by both improved information criterion and a significant F-test. Numbers within each quadrant indicate protein counts.
