## Supplementary material for "The human ovary exhibits dynamic molecular remodeling in the decades post-menopause": Suppl Figure S3

### Slide 1
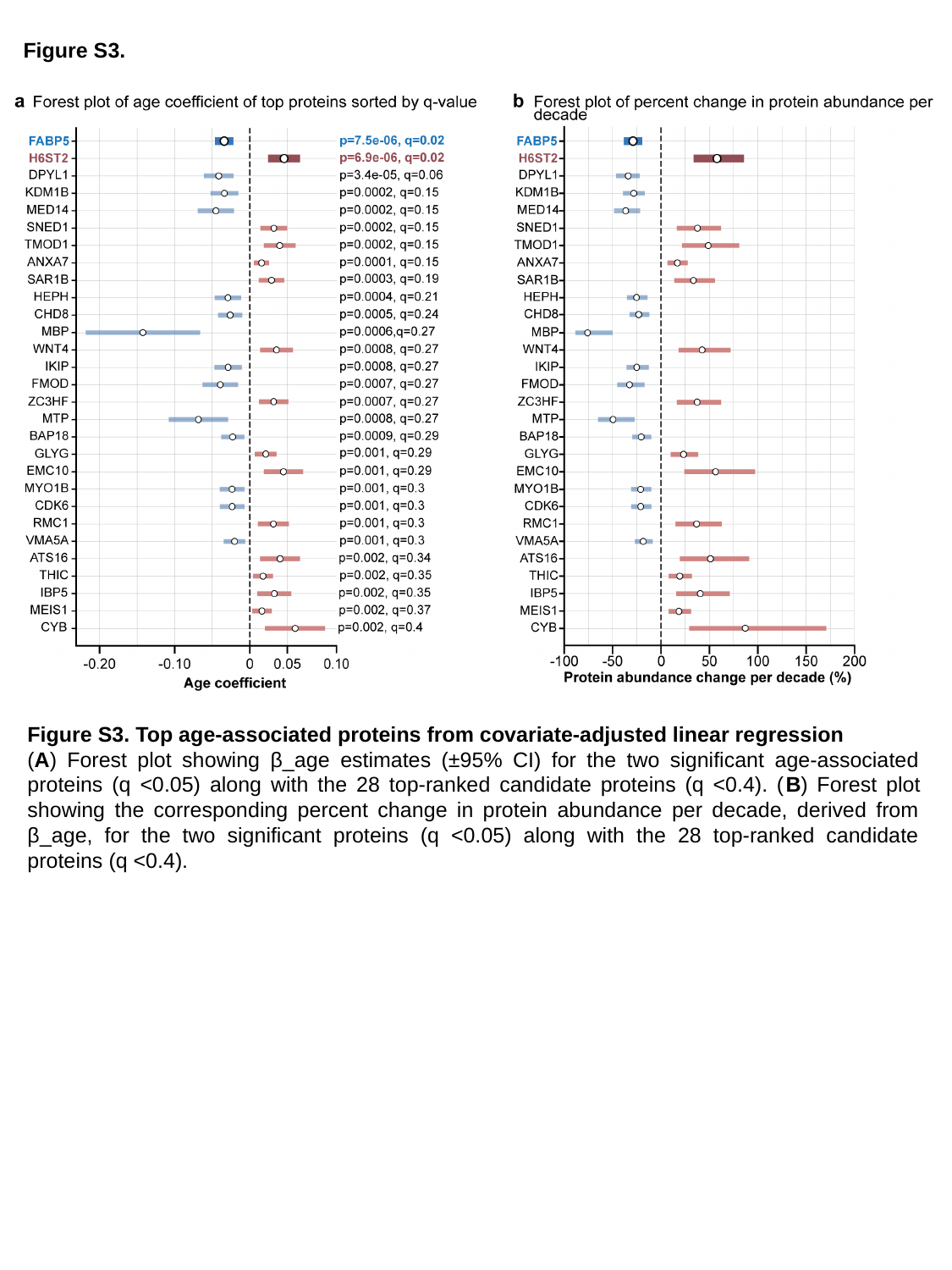

Figure S3.
Figure S3. Top age-associated proteins from covariate-adjusted linear regression
(A) Forest plot showing β_age estimates (±95% CI) for the two significant age-associated proteins (q <0.05) along with the 28 top-ranked candidate proteins (q <0.4). (B) Forest plot showing the corresponding percent change in protein abundance per decade, derived from β_age, for the two significant proteins (q <0.05) along with the 28 top-ranked candidate proteins (q <0.4).
