## Supplementary material for "The human ovary exhibits dynamic molecular remodeling in the decades post-menopause": Suppl Figure S4

### Slide 1
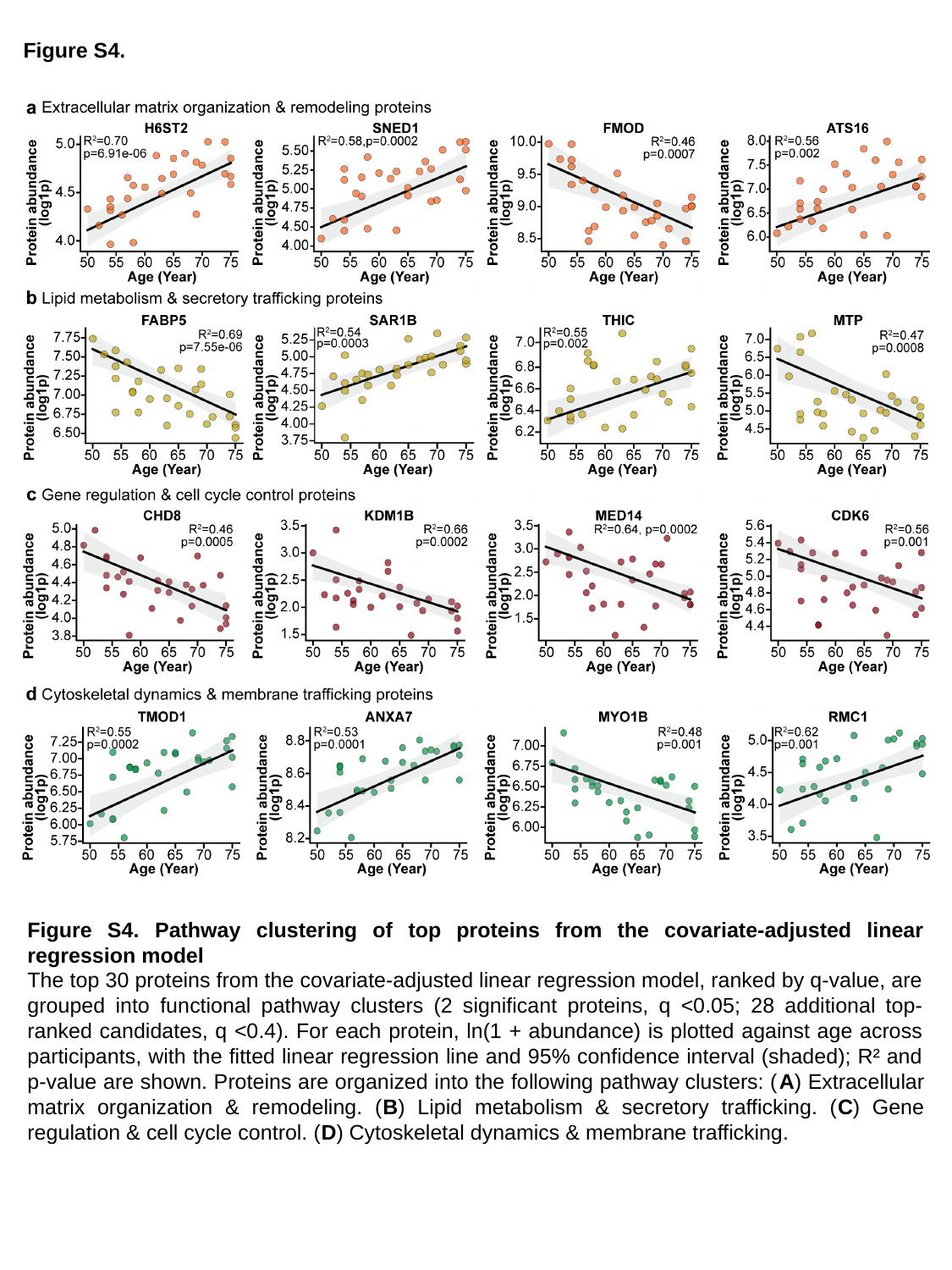

Figure S4.
Figure S4. Pathway clustering of top proteins from the covariate-adjusted linear regression model
The top 30 proteins from the covariate-adjusted linear regression model, ranked by q-value, are grouped into functional pathway clusters (2 significant proteins, q <0.05; 28 additional top-ranked candidates, q <0.4). For each protein, ln(1 + abundance) is plotted against age across participants, with the fitted linear regression line and 95% confidence interval (shaded); R² and p-value are shown. Proteins are organized into the following pathway clusters: (A) Extracellular matrix organization & remodeling. (B) Lipid metabolism & secretory trafficking. (C) Gene regulation & cell cycle control. (D) Cytoskeletal dynamics & membrane trafficking.
