## Supplementary material for "The human ovary exhibits dynamic molecular remodeling in the decades post-menopause": Suppl Figure S5

### Slide 1
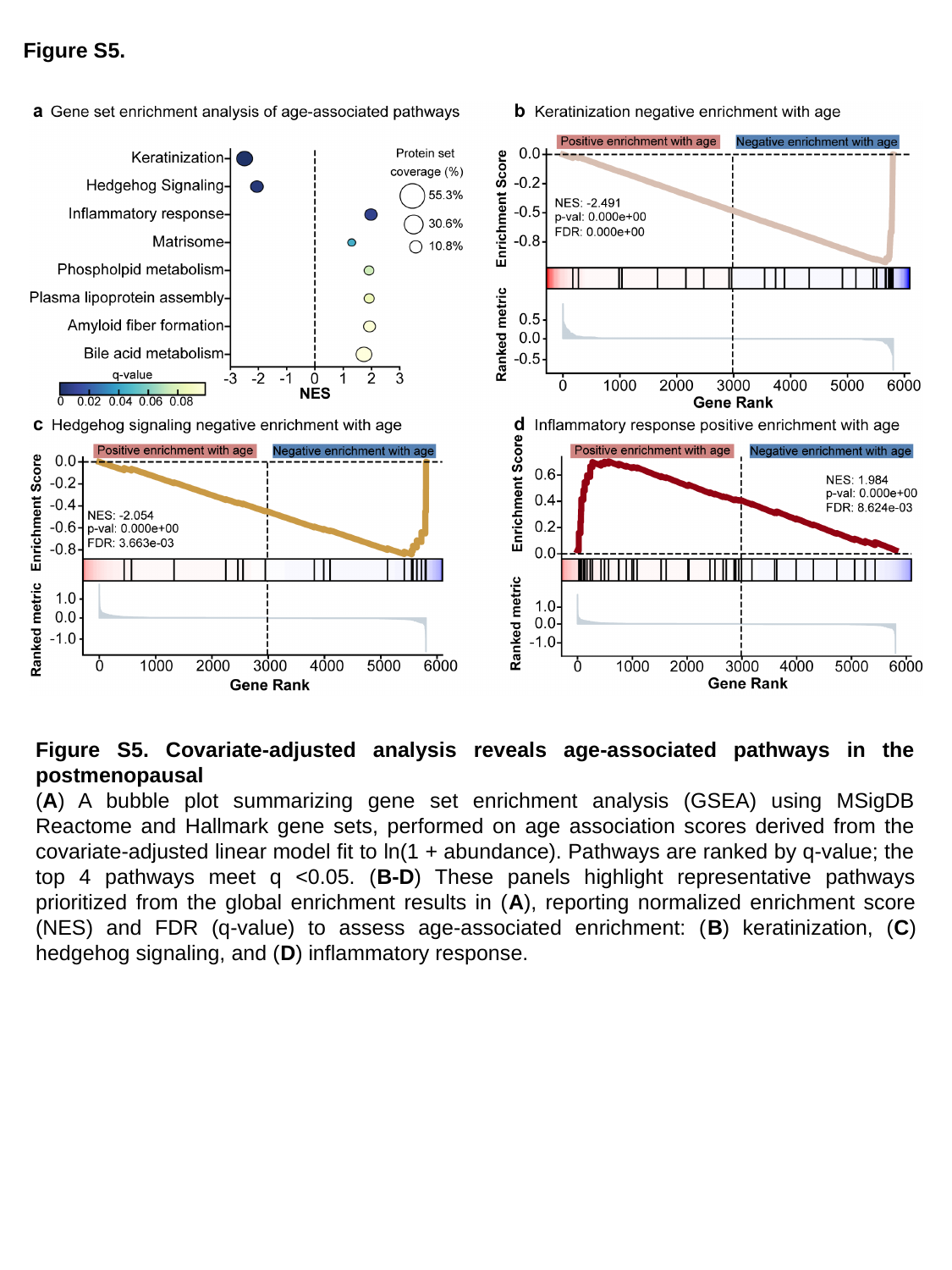

Figure S5.
Figure S5. Covariate-adjusted analysis reveals age-associated pathways in the postmenopausal
(A) A bubble plot summarizing gene set enrichment analysis (GSEA) using MSigDB Reactome and Hallmark gene sets, performed on age association scores derived from the covariate-adjusted linear model fit to ln(1 + abundance). Pathways are ranked by q-value; the top 4 pathways meet q <0.05. (B-D) These panels highlight representative pathways prioritized from the global enrichment results in (A), reporting normalized enrichment score (NES) and FDR (q-value) to assess age-associated enrichment: (B) keratinization, (C) hedgehog signaling, and (D) inflammatory response.
