## Supplementary material for "The human ovary exhibits dynamic molecular remodeling in the decades post-menopause": Suppl Figure S6

### Slide 1
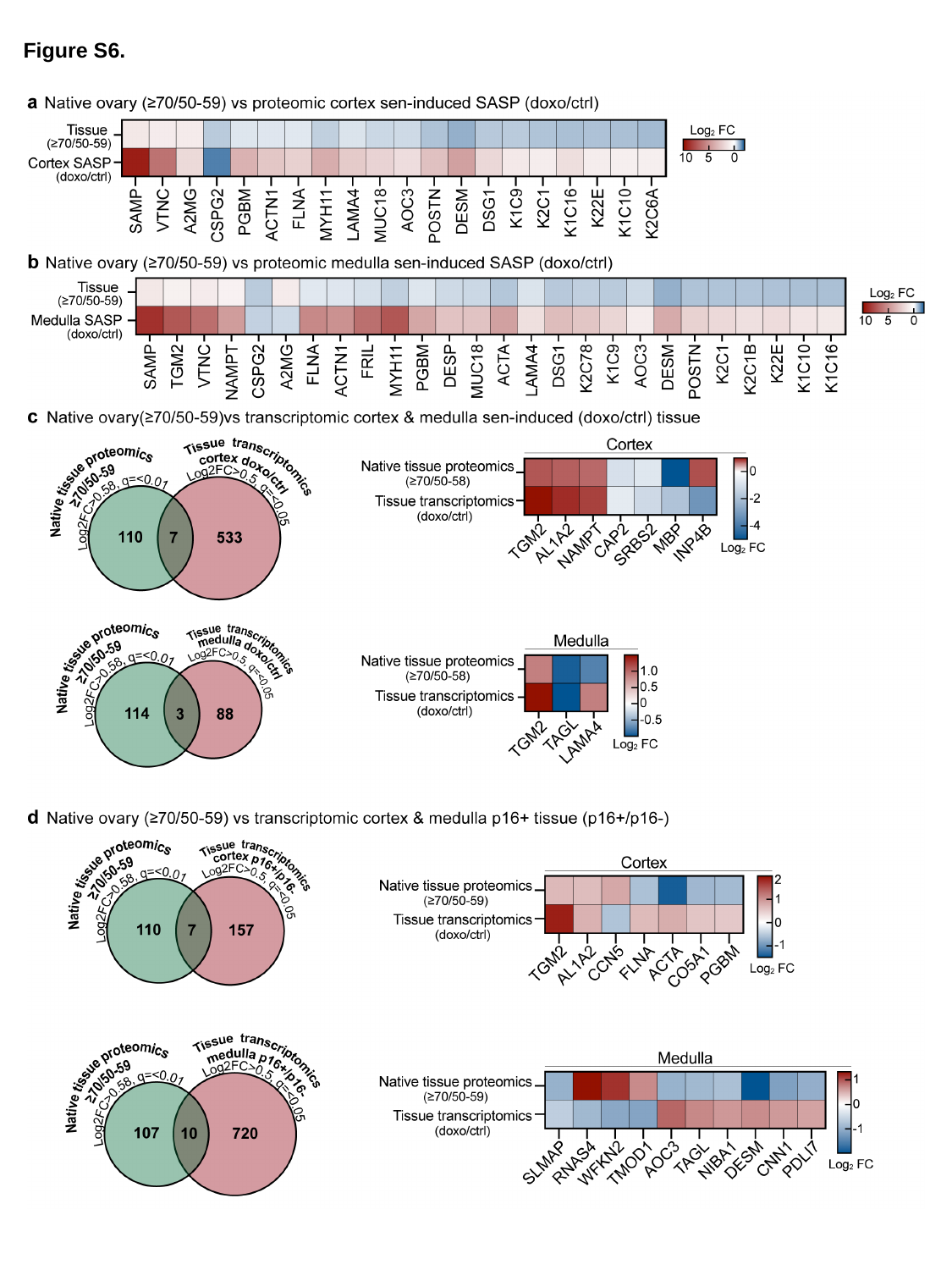

Figure S6.

### Slide 2
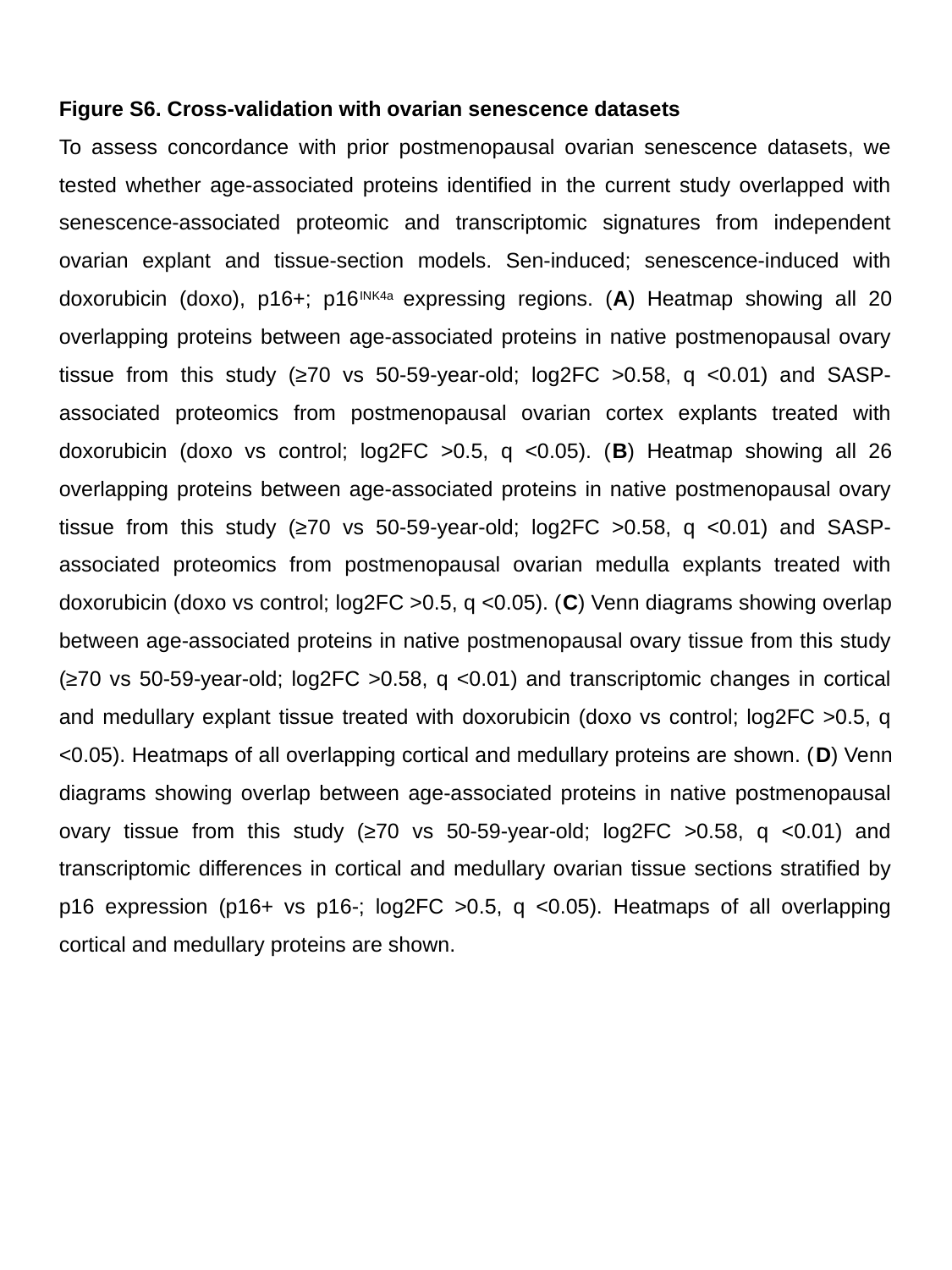

Figure S6. Cross-validation with ovarian senescence datasets
To assess concordance with prior postmenopausal ovarian senescence datasets, we tested whether age-associated proteins identified in the current study overlapped with senescence-associated proteomic and transcriptomic signatures from independent ovarian explant and tissue-section models. Sen-induced; senescence-induced with doxorubicin (doxo), p16+; p16^INK4a^ expressing regions. (A) Heatmap showing all 20 overlapping proteins between age-associated proteins in native postmenopausal ovary tissue from this study (≥70 vs 50-59-year-old; log2FC >0.58, q <0.01) and SASP-associated proteomics from postmenopausal ovarian cortex explants treated with doxorubicin (doxo vs control; log2FC >0.5, q <0.05). (B) Heatmap showing all 26 overlapping proteins between age-associated proteins in native postmenopausal ovary tissue from this study (≥70 vs 50-59-year-old; log2FC >0.58, q <0.01) and SASP-associated proteomics from postmenopausal ovarian medulla explants treated with doxorubicin (doxo vs control; log2FC >0.5, q <0.05). (C) Venn diagrams showing overlap between age-associated proteins in native postmenopausal ovary tissue from this study (≥70 vs 50-59-year-old; log2FC >0.58, q <0.01) and transcriptomic changes in cortical and medullary explant tissue treated with doxorubicin (doxo vs control; log2FC >0.5, q <0.05). Heatmaps of all overlapping cortical and medullary proteins are shown. (D) Venn diagrams showing overlap between age-associated proteins in native postmenopausal ovary tissue from this study (≥70 vs 50-59-year-old; log2FC >0.58, q <0.01) and transcriptomic differences in cortical and medullary ovarian tissue sections stratified by p16 expression (p16+ vs p16-; log2FC >0.5, q <0.05). Heatmaps of all overlapping cortical and medullary proteins are shown.
